# COVID-*e*Vax, an electroporated plasmid DNA vaccine candidate encoding the SARS-CoV-2 Receptor Binding Domain, elicits protective immune responses in animal models of COVID-19

**DOI:** 10.1101/2021.06.14.448343

**Authors:** Antonella Conforti, Emanuele Marra, Fabio Palombo, Giuseppe Roscilli, Micol Ravà, Valeria Fumagalli, Alessia Muzi, Mariano Maffei, Laura Luberto, Lucia Lione, Erika Salvatori, Mirco Compagnone, Eleonora Pinto, Emiliano Pavoni, Federica Bucci, Grazia Vitagliano, Daniela Stoppoloni, Maria Lucrezia Pacello, Manuela Cappelletti, Fabiana Fosca Ferrara, Emanuela D’Acunto, Valerio Chiarini, Roberto Arriga, Abraham Nyska, Pietro Di Lucia, Davide Marotta, Elisa Bono, Leonardo Giustini, Eleonora Sala, Chiara Perucchini, Jemma Paterson, Kathryn Ann Ryan, Amy-Rose Challis, Giulia Matusali, Francesca Colavita, Gianfranco Caselli, Elena Criscuolo, Nicola Clementi, Nicasio Mancini, Rüdiger Groß, Alina Seidel, Lukas Wettstein, Jan Münch, Lorena Donnici, Matteo Conti, Raffaele De Francesco, Mirela Kuka, Gennaro Ciliberto, Concetta Castilletti, Maria Rosaria Capobianchi, Giuseppe Ippolito, Luca G. Guidotti, Lucio Rovati, Matteo Iannacone, Luigi Aurisicchio

**Affiliations:** Takis Biotech, Via Castel Romano 100, 00128 Rome, Italy; Evvivax Biotech, Via Castel Romano 100, 00128 Rome, Italy; Neomatrix Biotech, Via Castel Romano 100, 00128 Rome, Italy; Division of Immunology, Transplantation and Infectious Diseases, IRCCS San Raffaele Scientific Institute, 20132 Milan, Italy; Sackler School of Medicine, Tel Aviv University, Haharuv 18, P.O. Box 184, Timrat, 36576, Israel; Vita-Salute San Raffaele University, 20132 Milan, Italy; National Infection Service, Public Health England (PHE), Porton Down, Salisbury, Wiltshire, United Kingdom, SP4 0JG; National Institute for Infectious Diseases Lazzaro Spallanzani, Via Portuense 292, 00149 Rome, Italy; Rottapharm Biotech s.r.l., Via Valosa di Sopra 9, 20900 Monza, Italy; Laboratory of Microbiology and Virology, IRCCS San Raffaele Scientific Institute, 20132 Milan, Italy; Institute of Molecular Virology, Ulm University Medical Center, Meyerhofstr. 1, 89081 Ulm, Germany; INGM - Istituto Nazionale di Genetica Molecolare “Romeo ed Erica Invernizzi”, Milan, Italy; National Cancer Institute Regina Elena, Via Elio Chianesi 53, 00144 Rome, Italy; Department of Pharmacological and Biomolecular Sciences, University of Milan, Italy; Experimental Imaging Centre, IRCCS San Raffaele Scientific Institute, 20132 Milan, Italy

**Keywords:** SARS-CoV-2, DNA vaccine, Antiviral Immunity, Animal models, Protection.

## Abstract

The COVID-19 pandemic caused by the β-coronavirus SARS-CoV-2 has made the development of safe and effective vaccines a critical global priority. To date, four vaccines have already been approved by European and American authorities for preventing COVID-19 but the development of additional vaccine platforms with improved supply and logistics profiles remains a pressing need. Here we report the preclinical evaluation of a novel COVID-19 vaccine candidate based on the electroporation of engineered, synthetic cDNA encoding a viral antigen in the skeletal muscle, a technology previously utilized for cancer vaccines. We constructed a set of prototype DNA vaccines expressing various forms of the SARS-CoV-2 Spike (S) protein and assessed their immunogenicity in animal models. Among them, COVID-*e*Vax – a DNA plasmid encoding a secreted monomeric form of SARS-CoV-2 S protein RBD – induced the most potent anti-SARS-CoV-2 neutralizing antibody responses (including against the current most common variants of concern) and a robust T cell response. Upon challenge with SARS-CoV-2, immunized K18-hACE2 transgenic mice showed reduced weight loss, improved pulmonary function and significantly lower viral replication in the lungs and brain. COVID-*e*Vax conferred significant protection to ferrets upon SARS-CoV-2 challenge. In summary, this study identifies COVID-*e*Vax as an ideal COVID-19 vaccine candidate suitable for clinical development. Accordingly, a combined phase I-II trial has recently started in Italy.

## Introduction

At the time of writing, SARS-CoV-2 has spread worldwide causing over 200 million confirmed cases and more than 4 million confirmed deaths.

To date, the regulatory agencies European Medicines Agency (EMA) and Food and Drug Administration (FDA) have authorized the conditional or emergency use of four vaccines against SARS-CoV-2, two based on mRNA (produced by Pfizer and Moderna) and two based on adenoviral vectors (produced by AstraZeneca and Johnson & Johnson). Additional vaccine candidates are under development and a continually updated list is available at https://www.who.int/publications/m/item/draft-landscape-of-covid-19-candidate-vaccines. Most COVID-19 vaccines and vaccine candidates target the SARS-CoV-2 full-length (FL) spike (S) glycoprotein, which mediates attachment and entry of the virus into host cells^1^, and employ both traditional and novel vaccine platforms such as inactivated virus, protein-based preparations, and virus-vectored and nucleic acid-based formulations^1, 2^.

Among the latter, DNA-based platforms show the greatest potential in terms of safety and ease of production^3^. Prior work has demonstrated that a DNA-based vaccine approach for SARS- and MERS-CoV induces neutralizing antibody (nAb) responses and provides protection in challenge models^4, 5^. Moreover, in a phase I dose-escalation study subjects immunized with a DNA vaccine encoding the MERS-CoV S protein showed durable nAb and T cell responses and a seroconversion rate of 96%^5^. The SARS-CoV-2 S protein is most similar in sequence and structure to SARS-CoV S and shares a global protein fold architecture with the MERS-CoV S protein^6^. Of note, the receptor-binding site of the S protein is a vulnerable target for antibodies. In fact, anti-MERS antibodies targeting the receptor-binding domain (RBD) of the S protein tend to have greater neutralizing potency than those directed to other epitopes^7^. More recently, a study by Piccoli *et al*.^8^ showed that depletion of anti-RBD antibodies in convalescent patient sera results in the loss of more than 90% neutralizing activity towards SARS-CoV-2, suggesting that the SARS-CoV-2 RBD represents a key target for vaccine development.

Here we describe the development of a DNA-based SARS-CoV-2 vaccine. Synthetic DNA is temperature-stable and cold-chain free, which are important advantages over approved RNA and vector vaccines for delivery to resource-limited settings. Furthermore, synthetic DNA vaccines are amenable to accelerated developmental timelines due to the relative simplicity by which multiple candidates can be designed, preclinically tested, manufactured in large quantities and progressed through established regulatory pathways to the clinic. Injection of DNA plasmid into the skeletal muscle followed by a short electrical stimulation – referred to as electro-gene-transfer or electroporation (EP) – enhances DNA uptake and gene expression by several hundred-fold^9–11^, leading to improved antigen expression and a local and transient tissue damage favoring inflammatory cell recruitment and cytokine production at the injection site^12^.

Exploiting our experience in the generation of vaccines based on the electroporation of plasmid DNA in the skeletal muscle^11^, we produced and screened several constructs expressing different portions of the SARS-CoV-2 S protein and identified COVID-*e*Vax – a DNA plasmid encoding a secreted monomeric form of SARS-CoV-2 S protein RBD – as a candidate for further clinical development. COVID-*e*Vax has a favorable safety profile, it induces potent anti-SARS-CoV-2 neutralizing antibody responses also against the current most common variants of concern (VOCs) as well as T cell responses, and it confers significant protection to hACE2 transgenic mice and ferrets upon SARS-CoV-2 challenge.

## Results

### DNA vaccine constructs and immunogenicity

We designed five different DNA constructs (Fig. 1A) encoding the following versions of the SARS-CoV-2 (Wuhan Hu-1, GenBank: MN908947) S protein: 1) the full-length protein (FL); 2) the receptor binding domain (RBD); 3) the highly variable N-terminal domain (NTD) and the RBD domain (N/R); 4) the whole S1 subunit (S1); 5) the RBD fused to a human IgG-Fc (RBD-Fc). To promote protein secretion, we introduced a tissue plasminogen activator (tPA) leader sequence in the RBD, N/R and S1 constructs and an IgK leader sequence in the RBD-Fc construct. Western blot analyses confirmed expression of all constructs in cell lysates and of S1, N/R and RBD in the culture supernatants (Fig. 1B).

**Figure 1.**
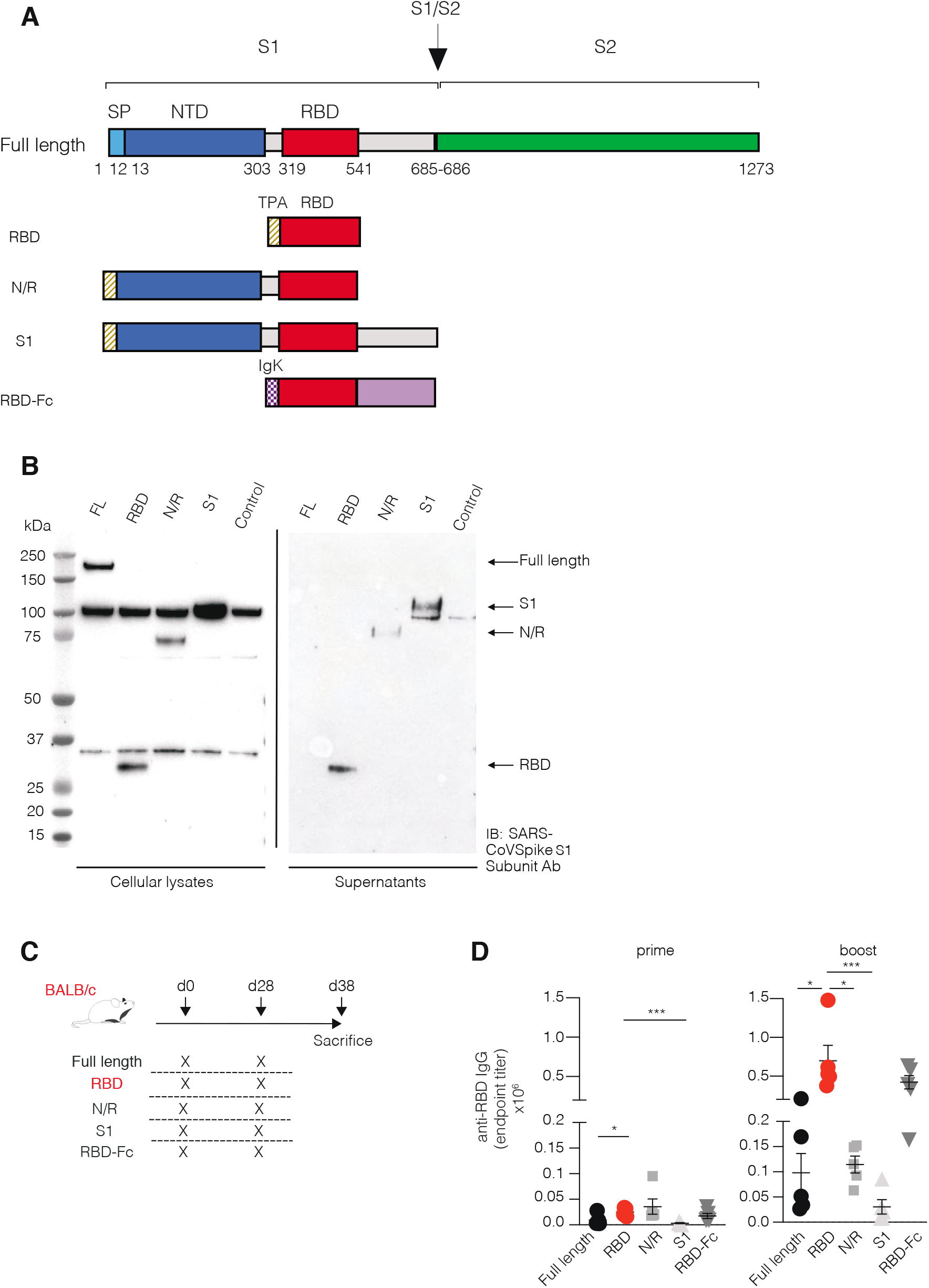
DNA vaccine constructs and immunogenicity. **(A)** Schematic representation of SARS-CoV-2 DNA vaccine construct candidates, encoding 1) the full-length protein (FL); 2) the receptor binding domain (RBD); 3) the highly variable N-terminal domain (NTD) and the RBD domain (N/R); 4) the whole S1 subunit (S1); 5) the RBD fused to a human IgG-Fc (RBD-Fc). The RBD, N/R and S1 constructs include a tPA leader sequence at the N-terminus, whereas the RBD-Fc construct contains a IgK leader sequence. **(B)** Western blot analysis of SARS-CoV-2 DNA vaccine constructs after transfection in HEK293 cells. Forty-eight hours after transfection, both cell lysates and supernatants were resolved on a gel and blotted with a polyclonal SARS-CoV Spike S1 Subunit antibody. Cells transfected with empty plasmid vector were used as negative control (control). Non-specific bands were detected both in cell lysates and in supernatants, likely due to non-specific binding of primary antibody. **(C)** Schematic representation of the experimental setup. Each DNA construct was injected intramuscularly (20 μg total, 10 μg each quadriceps) into BALB/c mice (*n* = 5) at day 0 (prime) and day 28 (boost). Intramuscular injection was followed by electroporation (EP). Mice were euthanized and analyzed at day 38. **(D)** Sera of BALB/c mice (*n* = 5) were collected at day 14 (only prime) and day 38 (prime-boost) and anti-RBD IgG levels were measured through ELISA, each dot represents a mouse. * p value < 0.05, *** p value < 0.001

Electroporation of these DNA vaccines in the skeletal muscle of BALB/c mice was adopted to evaluate immunogenicity. The vaccination protocol consisted of the injection of 20 μg of DNA into both quadriceps (10 μg of DNA in each muscle) into 6-week-old mice (Fig. 1C). A DNA plasmid expressing luciferase was used as a control for gene expression whereas a group of mice injected with DNA but not electroporated served as additional controls (Fig. S1). Mice received a second vaccination (boost) at day 28 and were sacrificed at day 38 (Fig. 1C). The humoral response in the sera of vaccinated mice was evaluated by measuring anti-RBD IgG titers by ELISA at day 14 (prime) and at day 38 (boost) (Fig. 1D). At day 14 all mice showed detectable anti-RBD IgG antibodies, and their levels significantly increased at day 38 (Fig. 1D). Notably, the most significant increase in antibody response was induced by the secreted RBD construct (Fig. 1D), with a calculated geometric mean of IgG endpoint titers^13^ as high as 1:24,223 after prime and 1:617,648 after boost. Since these preliminary data showed the RBD construct to be the most immunogenic among the five DNA constructs, RBD was chosen as the main vaccine candidate for further development and was directly compared with the FL construct in subsequent experiments.

### Detailed characterization of the humoral immune response elicited by the RBD vaccine candidate

We next sought to characterize the humoral response to the RBD vaccine in depth, focusing on the specificity, duration and neutralization capacity of the elicited antibodies. As per specificity, we carried out a B cell epitope mapping of the response elicited by the FL and RBD vaccines. To this end, a B cell ELISpot assay was performed by stimulating splenocytes collected from vaccinated BALB/c mice with 338 peptides covering the whole SARS-CoV-2 Spike protein. Sequences of positive hits (Table S1) were then mapped on the three-dimensional structure of the S protein^14^, hence outlining the epitope domains (Fig. 2A). Mice immunized with the RBD vaccine showed responses mapping mainly on conserved regions of the RBD, and not on regions most commonly affected by mutations in the current circulating VOCs (e.g., N501K, K417N, S477, E484K and L452, Fig. 2A). Despite the caveat that the abovementioned analysis detects linear and not conformational epitopes, this suggests that antibodies elicited by the RBD vaccine might be functional against the current most commonly circulating SARS-CoV-2 variants.

**Figure 2.**
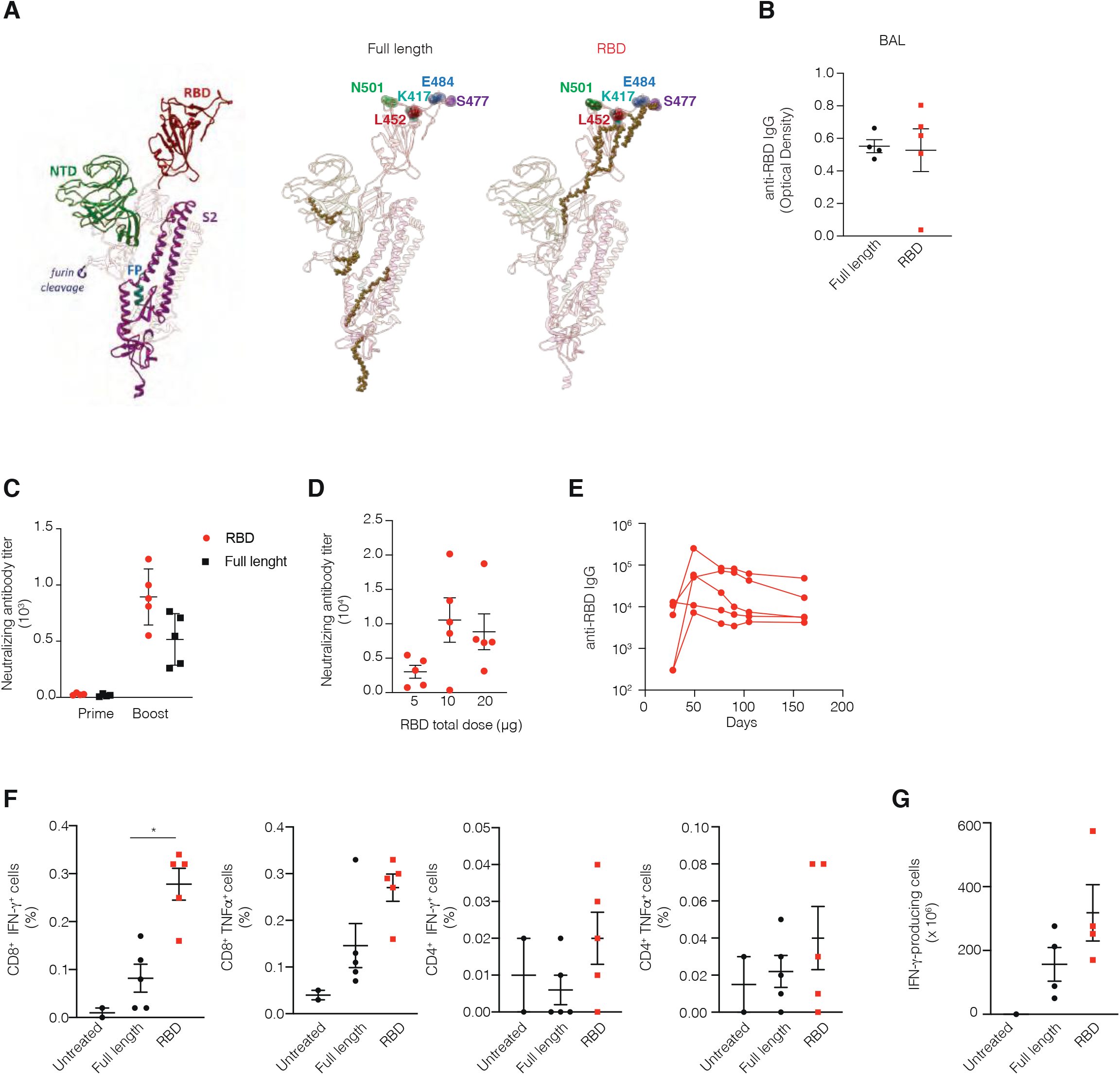
Characterization of the immune response elicited by the RBD vaccine candidate. **(A)** Antibody linear epitopes mapped onto the structure of the FL Spike protein. Each domain of the FL protein (NTD, RBD, furin cleavage, FP-fusion peptide and S2) is outlined with a different color (left panel) while the linear epitopes are shown as gold spheres within the Spike domains used for immunization (center and right panels). **(B)** Anti-RBD IgG levels measured in bronchoalveolar lavage (BAL) of FL and RBD-vaccinated BALB/c mice at day 38. **(C)** Neutralizing antibody titers in sera collected from RBD or FL-vaccinated BALB/c mice (*n* = 5) at day 14 (prime) and day 38 (boost), measured through a neutralization assay with infectious SARS-CoV-2-and Vero cells. **(D)** Neutralizing antibody titers in sera collected at day 38 from C57BL/6 (*n* = 5) vaccinated with increasing doses of RBD vaccine (5-10-20 μg) in a prime-boost regimen. **(E)** Serum anti-RBD IgG levels measured over time in sera of RBD-vaccinated C57BL/6 mice (prime-boost regimen, *n* = 5) up to 6 months starting from prime. **(F)** T cell immune response (IFN-γ^+^ and TNF-*α*^+^) in CD8^+^ and CD4^+^ cells measured by intracellular staining of splenocytes collected from FL- and RBD-vaccinated BALB/c mice (*n* = 5) at day 38 and restimulated with pool S1 peptides. **(G)** IFN-γ-producing T cells measured by ELISpot assay performed on BALs collected from BALB/c (*n* = 5) vaccinated with FL and RBD, intranasally challenged with 20 μg RBD protein at day 42 and culled the day after. * p value < 0.05

SARS-CoV-2-specific antibodies elicited by both the FL and the RBD vaccine were present not only in the sera but also in the lungs of vaccinated mice, as shown by analysis of bronchoalveolar lavages (BALs) of mice 38 days after vaccination (Fig. 2B).

We next sought to analyze whether the RBD-specific antibodies induced by the RBD vaccine were able to neutralize SARS-CoV-2. The neutralization capacity of antibodies induced by the RBD vaccine was comparable with FL after prime and superior after boost (Fig. 2C), with NT_50_ at day 38 of 894 ± 249. A dose-response experiment indicated that the neutralizing antibody titers plateaued at an RBD vaccine dose of 10 μg (injected in a single quadricep muscle) in a prime-boost regimen serum (Fig. 2D). Finally, total anti-RBD IgG antibodies persisted at high levels up to 6 months after vaccinations (Fig. 2E).

### Analysis of T cell responses elicited by the RBD vaccine candidate

Next we sought to evaluate the T cell response elicited by the RBD vaccine. To this end, we used peptide pools covering the S1 and S2 portions of the Spike protein (pools S1 and S2, respectively) to stimulate splenocytes collected from BALB/c mice at day 38 after vaccination (see Fig. 1C for experimental setup). IFN-γ released by T cells upon peptide re-stimulation was evaluated by ELISpot assay. As expected, in the group vaccinated with the RBD vaccine, we measured only T cell responses against pool S1 (that spans the RBD), whereas in the group vaccinated with the FL construct, we measured T cell responses against both pools S1 and S2 (data not shown). In order to reveal immunodominant epitopes eliciting the T cell response, we performed epitope mapping using thirty-seven matrix mapping pools, covering the entire sequence of the S protein (Fig. S2A), or twenty-four matrix pools, covering the RBD alone (Table S2 and Fig. S2B). Most of the H2-K^d^-restricted immunodominant epitopes were clustering in the RBD (Fig. S2C). Cytokine production by antigen-specific T cells was also evaluated by intracellular staining of splenocytes collected from vaccinated BALB/C mice and restimulated with pool S1 (Fig. 2F). Compared to FL, the RBD vaccine induced the highest frequency of CD8^+^ T cells producing either IFN-γ or TNF-*α* (Fig. 2F). To measure the potential recruitment of RBD-specific T-cells to the lungs, 20 μg of the RBD protein was injected intranasally in a group of vaccinated BALB/c mice two weeks after the second immunization and IFN-γ production from lymphocytes recovered from bronchoalveolar lavages (BAL) was measured by ELISpot assay one day later (Fig. 2G). Mice vaccinated with RBD showed a higher recruitment of RBD-specific T cells than mice vaccinated with the FL vaccine (Fig. 2H). A dose-response experiment conducted in C57BL/6 mice showed an even stronger specific T cell response than in BALB/c mice (data not shown) and a clear dose-dependency (Fig. S3A). A nonlinear fitting analysis of the curve (after pool S1 stimulation) revealed an ED50 of 2.06 ± 0.86 μg (Fig. S3B). Cytokine analyses in vaccinated C57BL/6 mice revealed a predominant IFN-γ- and TNF-*α*-producing CD8^+^ T cell response, independently of the sex and age of the mice (Fig. S3C and data not shown).

### Safety and immunogenicity of the RBD vaccine candidate in rats

We next evaluated the safety and immunogenicity of the RBD vaccine candidate in rats, an animal model highly suited for toxicological studies. Seven-week-old female Sprague-Dawley rats were injected intramuscularly with PBS or 100, 200 or 400 μg of the RBD vaccine (divided equally in the two quadriceps) followed by EP at day 0 and day 14 (Fig. 3A). The immunizations were well tolerated, with only mild to moderate lesions^15^ at the injection site that were almost fully recovered within four weeks (Fig. S4), and an increased cellularity in the draining lymph nodes (data not shown).

**Figure 3.**
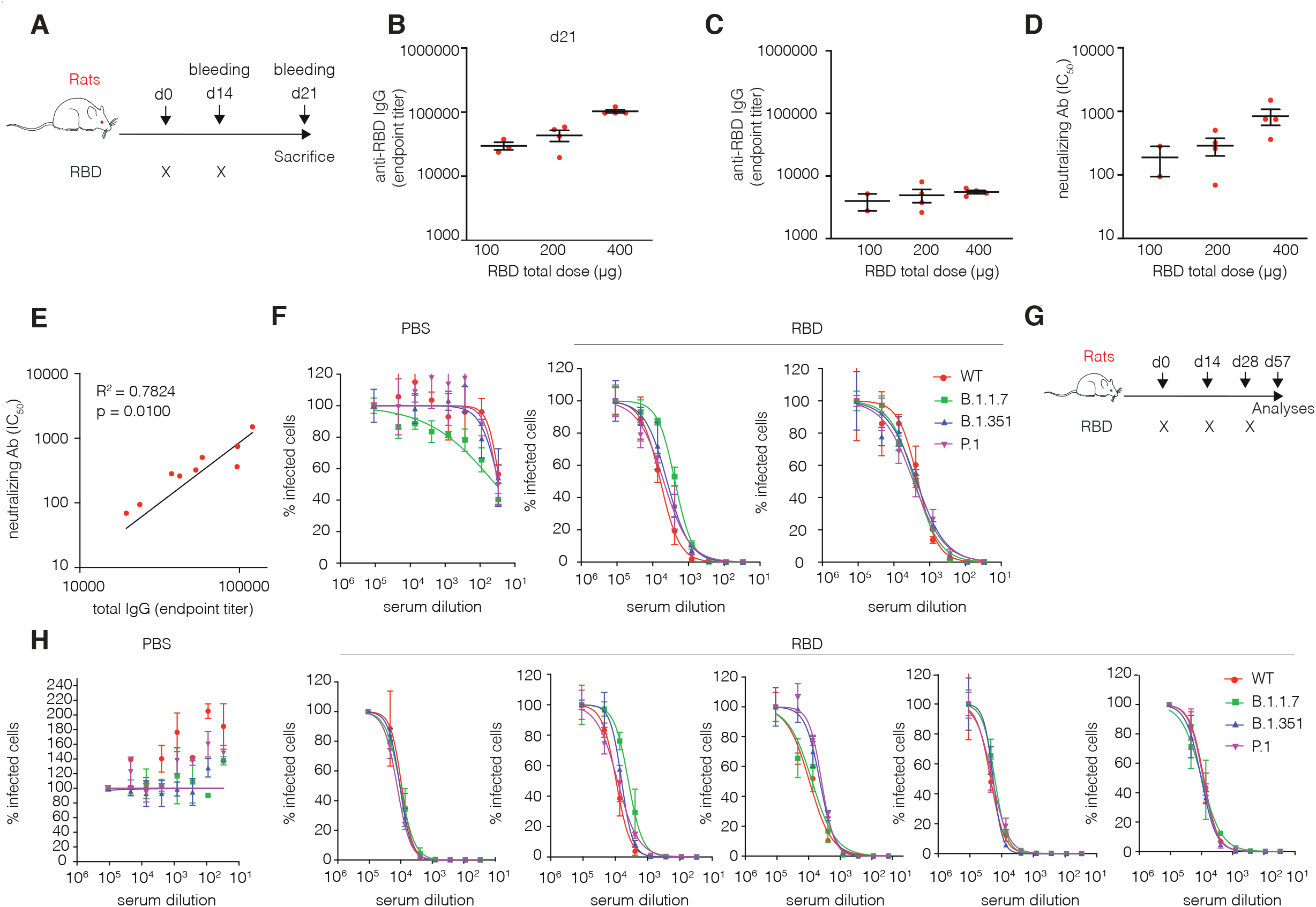
Immunogenicity of the RBD vaccine in rats. **(A)** Schematic representation of the experimental setup. Sprague-Dawley rats (*n* = 16) received two doses of RBD vaccine (day 0 and day 14) via intramuscular injection followed by EP. **(B)** Total IgG endpoint titer measured by ELISA assay performed on sera collected at day 14 (prime) from rats vaccinated with increasing doses of RBD (100-200-400 μg). **(C)** Total IgG endpoint titer measured by ELISA assay performed on sera collected at day 21 (prime-boost) from rats vaccinated with increasing doses of RBD vaccine. **(D)** Neutralizing antibody titer (IC_50_) of sera collected from the same rats of panel C. **(E)** Correlation between total IgG endpoint titers and neutralizing antibody IC_50_ values. **(F)** Dose response curve representing neutralization activity of plasma against SARS-CoV-2 pseudovirus carrying the SPIKE protein of wild type (WT) virus or variants (B.1.1.7, B.1.351 and P.1). Plasma was collected at sacrifice from rats vaccinated with 400 μg of the RBD vaccine (two dose vaccination regimen, day 0 and day 14) or with PBS, as negative control **(G)** Schematic representation of the experimental setup. Five Sprague-Dawley rats received three doses of RBD vaccine (day 0, 14 and 28) via intramuscular injection followed by EP. **(H)** Dose response curve representing neutralization activity of plasma against SARS-CoV-2 pseudovirus carrying the SPIKE protein of wild type (WT) virus or variants (B.1.1.7, B.1.351 and P.1). Plasma was collected at sacrifice from rats vaccinated with 400 μg of the RBD vaccine (three dose vaccination regimen, day 0, 14 and 28) or with PBS, as negative control.

Quantification of the RBD-specific antibody titers showed a robust and dose-dependent antibody production, with ELISA endpoint titers up to 152,991 for the highest dose (Fig. 3B, C). Immunization with the RBD vaccine induced high neutralizing antibody titers (Fig. 3D) which correlated with the total IgG endpoint titers (Fig. 3E). Finally, sera from Sprague-Dawley rats that were immunized with 400 μg of the RBD vaccine at day 0 and 14, or that received a third dose at day 28, were assessed for neutralizing activity against three major SARS-CoV-2 variants (i.e., B.1.1.7, B.1.351 and P.1) utilizing a lentiviral pseudotyped assay (Fig. 3F-H).

### The RBD vaccine candidate elicits protective immune responses in K18-hACE2 transgenic mice and in ferrets

To explore the *in vivo* protection efficacy of RBD vaccine against SARS-CoV-2 challenge, K18-hACE2 transgenic mice^16^ received two intramuscular immunizations (at day -39 and at day -18) of 10 μg of the RBD vaccine (*n* = 7) or PBS (*n* = 6) followed by EP (Fig. 4A). Pre-challenge sera collected 1 day prior to SARS-CoV-2 infection showed that the RBD vaccine induced robust RBD-specific IgG antibodies (average concentration of ∼ 50 μg/ml, Fig. 4B). Eighteen days after the boost immunization, all mice were infected intranasally with 1 × 10^5^ TCID_50_ of SARS-CoV-2 (hCoV-19/Italy/LOM-UniSR-1/2020; GISAID Accession ID: EPI_ISL_413489) (Fig. 4A). As expected^17^, beginning 3-4 days post infection (p.i.) PBS-treated K18-hACE2 transgenic mice infected with SARS-CoV-2 exhibited a weight loss close to 20% of their body weight and a lethargic behavior (Fig. 4C and data not shown). By contrast, K18-hACE2 transgenic mice immunized with RBD vaccine maintained stable body weight upon SARS-CoV-2 challenge and appeared more active (Fig. 4C and data not shown). We used whole-body plethysmography to evaluate several complementary metrics of pulmonary function, obstruction, and bronchoconstriction, including frequency, enhanced pause (PenH), and the fraction of expiration time at which the peak occurs (Rpef)^18, 19^. PBS-treated mice infected with SARS-CoV-2 exhibited a decreased respiratory rate (Fig. 4D), an increased PenH (Fig. 4E) and a decreased Rpef (Fig. 4F), indicative of pronounced loss of pulmonary function. By contrast, K18-hACE2 transgenic mice immunized with the RBD vaccine prior to infection maintained a relatively stable respiratory rate (Fig. 4D), had a much lower PenH (Fig. 4E) and a higher Rpef (Fig. 4F), indicative of better pulmonary function. Much higher amounts of viral RNA, infectious SARS-CoV-2 and viral N protein were detected in the lungs and brain of PBS-treated mice compared to mice immunized with the RBD vaccine (Fig. 4G-L). The lower viral titers in the lungs of immunized mice were associated with the detection of RBD-specific CD4^+^ T cells producing IFN-γ, TNF-*α* or both as well as RBD-specific IFN-γ-producing CD8^+^ T cells (Fig. 4M).

**Figure 4.**
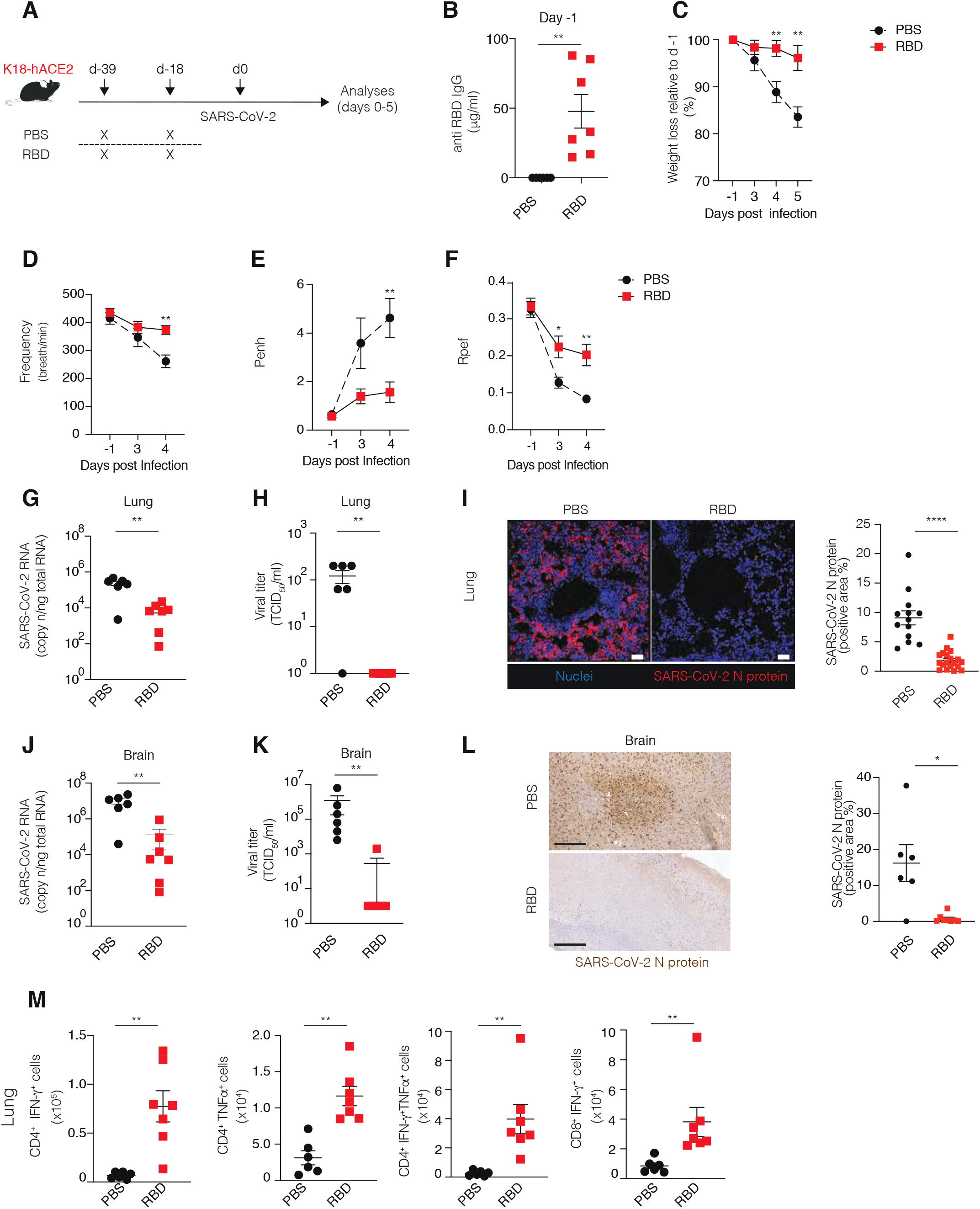
In vivo protection efficacy of RBD vaccine against SARS-CoV-2 virus challenge in hACE2 transgenic mice. **(A)** Schematic representation of the experimental setup. K18-hACE2 (C57BL/6) mice received two immunizations (day -39, day -18) of 10 μg RBD vaccine (*n* = 7) or PBS (*n* = 6) via intramuscular injection followed by EP before intranasal challenge with SARS-CoV-2. Lung and brain were collected and analyzed five days after SARS-CoV-2 infection. **(B)** Serum anti-RBD IgG levels of RBD vaccine or PBS challenged mice detected by ELISA assay; sera were collected right before SARS-CoV-2 infection. **(C)** Mouse body weights were monitored daily for up to 5 days. PBS-treated mice showed a rapid body weight decrease from day 4, instead RBD vaccine-challenged mice demonstrated normal statuses. **(D-F)** Whole-body plethysmography assessing pulmonary function for Frequency **(D)**, Penh **(E)** and Rpef **(F)**. **(G)** SARS-CoV-2 RNA in the lung was quantified by RT–qPCR 5 days after infection. **(H)** Viral titers in the lung 5 days after infection were determined by median tissue culture infectious dose (TCID_50_). **(I)** Representative confocal immunofluorescence micrographs of lung sections from PBS-treated mice (left) or RBD-treated mice (right) 5 days after SARS-CoV-2 infection. N-SARS-CoV-2 positive cells are depicted in red and nuclei in blue. Scale bars represent 30 μm. Right panel, quantification of N-SARS-CoV-2 signal, each dot represents a different section. **(J)** SARS-CoV-2 RNA in the brain was quantified by quantitative PCR with reverse transcription (RT–qPCR) 5 days after infection. **(K)** Viral titers in the brain 5 days after infection were determined by median tissue culture infectious dose (TCID_50_). **(L)** Representative immunohistochemical micrographs of brain sections from PBS-treated mice (top) or RBD-treated mice (bottom) 5 days after SARS-CoV-2 infection. N-SARS-CoV-2 expression is shown in brown. Scale bars, 300 μm. Right panel, quantification of N-SARS-CoV-2 signal, each dot represents a mouse. **(M)** Absolute numbers of CD4^+^ T cells producing IFN-γ, TNF-*α* or both and of CD8^+^ T cells producing IFN-γ in the lung of the indicated mice five days after SARS-CoV-2 infection. * p value < 0.05, ** p value < 0.01, **** p value < 0.0001

Besides inducing potent adaptive immune responses, the protection induced by the RBD vaccine might lie in the competitive inhibition of SARS-CoV-2 binding to ACE2 by the secreted RBD. Indeed, RBD is detectable in the sera and in the BAL of immunized BALB/c mice as early as 2 days after immunization (Fig. S5A-C), time point in which anti-RBD antibodies are not yet detectable (Fig. S5D). To test whether this secreted RBD would compete with SARS-CoV-2 for ACE2 binding, we immunized K18-hACE2 transgenic mice with the RBD vaccine 2 days prior to intranasal inoculation with a luciferase-encoding lentiviral vector pseudotyped with the SARS-CoV-2 Spike protein. Two days later the lungs of treated mice were assessed for bioluminescence using an *in vivo* imaging system (IVIS). As shown in Fig. S5E, compared to mice injected with PBS, mice immunized with the RBD vaccine exhibited a reduced bioluminescence, indicative of a significantly lower *in vivo* transduction. Further experiments should determine the extent to which the abovementioned mechanism confers protection by the RBD vaccine.

To confirm the immunogenicity and protective efficacy of the RBD vaccine against SARS-CoV-2 infection in a different and larger animal model, sixteen female ferrets weighing over 750 g were either left untreated (control) or injected with 400 μg of the RBD vaccine followed by electroporation 42 and 14 days prior to intranasal infection with 5 x 10^6^ pfu of SARS-CoV-2 isolate Victoria/1/2020 (Fig. 5A). Compared to control animals, viral subgenomic RNA detected in nasal washes and throat swabs at day 7 post challenge in immunized ferrets were significantly reduced (Fig. 5B, C).

**Figure 5.**
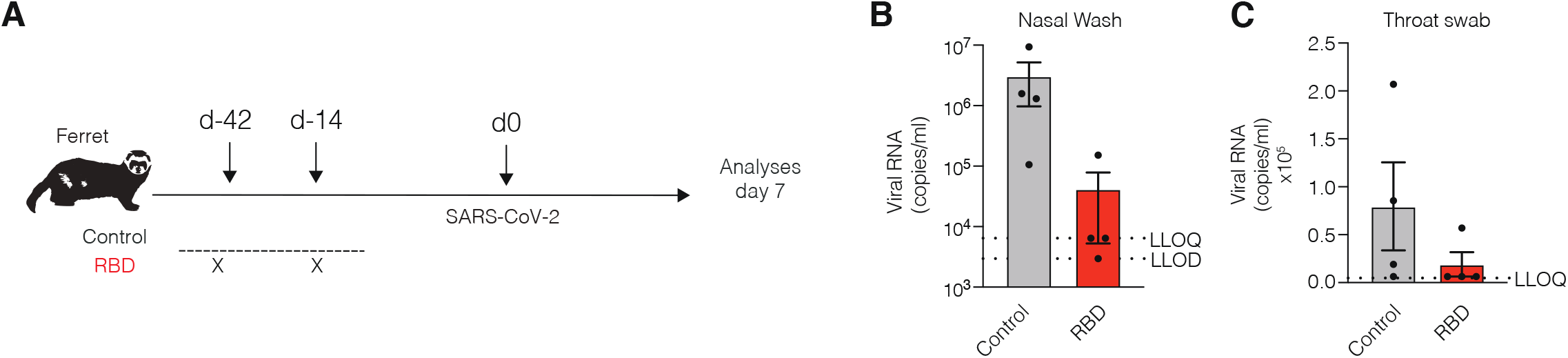
Evaluation of RBD vaccine efficacy in a ferret infection model. **(A)** Schematic representation of the experimental setup. Female ferrets (*n* = 8) were either left untreated (control) or received two immunizations (day -42, day -14) of 400 μg RBD vaccine via intramuscular injection followed by electroporation before intranasal challenge with 5×10^6^ PFU/ml of SARS-CoV-2. Four animals from each group were euthanized at each timepoint (3 and 7 days post challenge). **(B)** Viral RNA detected in nasal wash from the control group or the vaccinated group following challenge. Results below the lower limit of detection (LLOD) have been assigned a value of 1157 copies/ml, and results between the LLOD and lower limit of quantification (LLOQ) have been assigned a value of 6429 copies/ml. **(C)** Viral RNA detected in throat swabs from the control group or the vaccinated group following challenge. * p value < 0.05

Together, the results obtained in two distinct animal models of SARS-CoV-2 infection indicate that the RBD vaccine induces protective immune responses.

## Discussion

COVID-19 pandemic has stimulated the evaluation and the approval of novel genetic vaccination platform technologies in a very short time frame. In fact, platforms based on Adenoviral vectors and mRNA vaccines had been evaluated only in clinical trials but never approved as products. COVID-*e*Vax is based on DNA. Thus far, DNA vaccines have been approved only for veterinary applications and are being evaluated in Oncology up to Phase III trial. A drawback of DNA-based vaccines has been the poor immunogenicity when moving from mice to larger species. However, the use of electroporation and other delivery technologies has greatly enhanced their potency. Similar to other DNA-based vaccines, such as those utilizing adenoviral vectors, the need for nuclear delivery, the risk for chromosomal integration, the potential activation of oncogenes and induction of anti-DNA antibodies needs to be taken into account. This safety concerns are carefully explored according to indications provided by the regulatory agencies (FDA Guidance for Industry - Considerations for Plasmid DNA Vaccines for Infectious Disease Indications. U.S. Department of Health and Human Services, Food and Drug Administration, Center for Biologics Evaluation and Research; November 2007) and WHO (https://www.who.int/teams/health-product-policy-and-standards/standards-and-specifications/vaccines-quality/dna). Very recently (August 21, 2021), a DNA vaccine (ZyCoV-D) against SARS-CoV-2 developed by Zydus/Cadila has achieved positive results in a Phase III clinical trial and was approved for emergency use authorization (EUA) with the office of the Drug Controller General of India (DCGI). We believe that this vaccine will pave the way for the approval of other DNA vaccines, such as COVID-*e*Vax.

DNA-based vaccines are engineered for maximal gene expression and immunogenicity, they can be quickly designed from new genetic viral sequences, and they allow for fast and scalable manufacturing as well as long-term stability at room temperature. Moreover, DNA vaccines do not require complex formulations such as those based on nanoparticles (necessary for peptide- or RNA-based vaccines). An efficient DNA uptake can be obtained with different methods^20^. Among others, electroporation (EP) increases the initial uptake of DNA plasmid by local cells by approximately 500-fold^20–22^. Here, we adopted an EP technology manufactured by the Italian company IGEA, a leader in tissue EP, and extensively tested both in mice and other animal species^11, 23–26^. This platform technology has been referred to as X-*e*Vax, where X represents the antigen (or the disease).

Multiple studies have reported that DNA vaccines allow for the generation of cellular and humoral responses against pathogens, making this platform ideal for rapid vaccine development against emerging infectious diseases^27^. Among these, three DNA vaccines targeting coronavirus S protein have already been tested in humans. The first candidate DNA vaccine expressing SARS-CoV S protein was VRC-SRSDNS015-00-VP and was tested in 10 healthy adults, aged 21 to 49 years, in 2004 and 2005^28^. This vaccine was administered intramuscularly at a dose of 4 mg by a Biojector needle-free device and proved to be safe and immunogenic. Another candidate DNA vaccine expressing MERS S protein was GLS-5300, evaluated in 75 healthy subjects, aged 19 to 50 years, in 2016^5^. GLS-5300 DNA vaccine was also administered intramuscularly followed by EP, in a dose escalation trial at 0.67, 2 or 6 mg. Overall, the vaccine was safe and immunogenic, as assessed by seroconversion and vaccine-induced T cell responses in most vaccine recipients^5^. The third candidate DNA vaccine, INO-4800, expressing the SARS-CoV-2 S protein, has been evaluated in 40 healthy subjects, aged 18 to 50 years, at doses of 1 or 2 mg. Administration by intradermal injection followed by EP using Inovio’s Cellectra^®^ device has proven generally safe and immunogenic as assessed by humoral and cellular immune responses^29^. INO-4800 is currently investigated in a Phase 2/3 clinical trial (NCT04642638).

The RBD vaccine candidate reported here, referred to hereafter as COVID-*e*Vax, was selected among 5 different candidates, encoding either the full-length S protein, or portions of it or engineered versions. Preclinical results have shown that all versions were capable of generating antibodies against the RBD region, key for viral entry. The RBD vaccine was chosen over the other candidates not only for its capacity to elicit potent neutralizing antibodies, but also in light of the capacity to induce a robust T cell response, the observation that anti-RBD antibodies represent >90% of the neutralizing antibodies in convalescent patients^8, 30^ and the notion that an RBD vaccine might be devoid of potential antibody-dependent enhancement. Moreover, a recent report suggests that RBD vaccines could better promote the elicitation of high titers of broad sarbecovirus neutralizing antibodies owing to enhanced accessibility of appropriate antigenic sites compared to the current full length vaccines^31^. Safety and immunogenicity of COVID-*e*Vax was demonstrated in mice and rats. Antibodies binding to the RBD domain of SARS-CoV-2 were detected not only in sera but also in the lungs of vaccinated mice and their functionality was assessed through neutralization of both wild-type virus and pseudovirus and by competitive inhibition of SARS-CoV-2 S protein binding to the ACE2 receptor in the presence of sera bled from immunized mice (data not shown). Besides the humoral response, a consistent T cell response, thought to contribute to preventing severe forms of COVID-19 in humans^32^, was elicited by a prime-boost vaccination schedule of COVID-*e*Vax. Moreover, since COVID-*e*Vax targets a small region within the S protein, i.e. the RBD, the risk of inducing an antibody-dependent enhancement (ADE) should be minimal.

The protective role played by the anti-RBD humoral and T cell response was demonstrated in both K18-hACE2 transgenic mice as well as in ferrets. Besides inducing potent adaptive immune responses, a potential additional mechanism of action of COVID-*e*Vax might be due to the competitive inhibition of SARS-CoV-2 binding to ACE2 by the secreted RBD. This is an intriguing potential mechanism that awaits confirmation and further characterization.

COVID-*e*Vax is currently being evaluated in the COV-1/2-01 Phase I/II study (EudraCT 2020-003734-20) at four clinical sites in Italy. The trial is aimed at assessing the safety and immunogenicity of COVID-*e*Vax in healthy subjects of both genders, 18 to 65 years of age. In the Phase I Dose Escalation part, COVID-*e*Vax is administered at 3 escalating doses (20 subjects/cohort), in a prime-boost setting (4 weeks apart), from 0.5 to 2 mg/dose. In addition, a cohort in a single 2 mg dose schedule will also be tested. The vaccine is administered by the intramuscular route followed by EP using the IGEA new ElectroPoration System (EPSGun), and the EGT technology for pulse generation (Cliniporator®) commercially available in the EU. In both phases, subjects will be followed up for a total duration of 6 months after the first vaccination.

In summary, COVID-*e*Vax is a highly efficient vaccination platform capable of inducing robust, protective neutralizing antibody and T cell responses in a variety of animal models. COVID-*e*Vax can be administered multiple times, without the risk of inducing antibody responses to the vaccine itself, which may happen in the case of virus-based vector vaccines. We believe that the DNA vaccination platform described here offers unique advantages over other candidate vaccines, such as rapid manufacturing in response to sequence mutations (compared to protein- or viral vector-based vaccines), and greater stability at room temperature (compared to RNA-based platforms). With an increasing number of people having been immunized against SARS-CoV-2 with an RNA-, adenovirus-, or protein-based vaccine, COVID-*e*Vax might be also considered as an additional platform for booster immunizations to extend the duration of protective immunity.

## Acknowledgments

We thank the entire Rottapharm Biotech and IGEA Teams for useful scientific and regulatory discussions and for setting up the EPSgun technology in a short time. We also thank M. Mainetti, M. Freschi, A. Fiocchi for technical support; M. Silva for secretarial assistance; and the members of the Iannacone laboratory for helpful discussions. Flow cytometry was carried out at FRACTAL, a flow cytometry resource and advanced cytometry technical applications laboratory established by the San Raffaele Scientific Institute. Confocal immunofluorescence histology was carried out at Alembic, an advanced microscopy laboratory established by the San Raffaele Scientific Institute and the Vita-Salute San Raffaele University. We would like to acknowledge the PhD program in Basic and Applied Immunology and Oncology at Vita-Salute San Raffaele University, as D.M. and E.S. conducted this study as partial fulfillment of their PhD in Molecular Medicine within that program. M.I. is supported by the European Research Council (ERC) Consolidator Grant 725038, ERC Proof of Concept Grant 957502, Italian Association for Cancer Research (AIRC) Grants 19891 and 22737, Italian Ministry of Health Grants RF-2018-12365801 and COVID-2020-12371617, Lombardy Foundation for Biomedical Research (FRRB) Grant 2015-0010, the European Molecular Biology Organization Young Investigator Program, and Funded Research Agreements from Gilead Sciences, Takis Biotech, Toscana Life Sciences and Asher Bio. L.G.G. is supported by the Italian Association for Cancer Research (AIRC) Grant 22737, Lombardy Open Innovation Grant 229452, PRIN Grant 2017MPCWPY from the Italian Ministry of Education, University and Research, Funded Research Agreements from Gilead Sciences, Avalia Therapeutics and CNCCS SCARL and donations from FONDAZIONE SAME and FONDAZIONE PROSSIMO MIO for COVID-19-related research. M.K. is supported by the Italian Ministry of Education, University and Research grant PRIN-2017ZXT5WR. J.M. is supported by funding from the Deutsche Forschungsgemeinschaft through Fokus-Förderung COVID-19 and the CRC1279, the European Union Horizon 2020 Framework Programme for Innovation and Research (Fight-nCoV), the Ministry for Science, Research and Arts of Baden-Württemberg, and the BMWi–Federal Ministry for Economic Affairs and Energy (COMBI-COV-2). R.G., A.S., and L.W. are part of the International Graduate School in Molecular Medicine Ulm. Takis Research activities are supported in part by the Italian Ministry of Economic Development through grants F/050298/02/X32, F/090033/01-04/X36 and F/190180/01/X44. We are also grateful to Lazio Innova for the funding provided through the grant A0376-2020-0700050 Prog. T0002E0001 “Emergenza Coronavirus ed oltre”, to Vitares no profit organization and Fondazione Melanoma for providing support to develop the assays used in clinical trials. PHE activities (Covivax project) are supported by the European Network of vaccine research and development (TRANSVAC2).

## Author contributions

A.C., E.M., F.P., G.R., M.R., V.F., A.M., M.M., L. Luberto, L. Lione, E. Salvatori, M.C., E. Pinto, E. Pavoni, F.B., G.V., D.S., M.L.P., M.C., F.F.F., E.D., V.C, A.N., P.D.L., D.M., E.B., L.G., E. Sala, C.P., J.P., K.A.R., A.R.C., G.M., G.C., E.C., N.C., N.M., R.G., A.S., L.W., L.D., M.C. performed experiments; A.C., M.R., V.F., E.S., G.C., J.M., L.D., M.C., R.D.F. analyzed and interpreted data; A.C. and M.R. prepared the figures; J.M., R.D.F. M.K., G.C., C.C., M.R.C., G.I., L.G.G., L.R., provided funding, conceptual advice and edited the manuscript; M.I. and L.A. coordinated the study, provided funding and wrote the paper.

## Data and materials availability

All data are available in the main text or the supplementary material.

## Materials and Methods

### Synthetic genes and constructs

The synthesis and codon optimization analysis of a cDNA encoding the SARS-CoV-2 protein S has been performed at Genscript (China). All constructs were completely synthetic and optimized for codon usage. Codon-optimized variants took into account codon usage bias, GC content, CpG dinucleotides content, mRNA secondary structure, cryptic splicing sites, premature PolyA sites, internal chi sites and ribosomal binding sites, negative CpG islands, RNA instability motif (ARE), repeat sequences (direct repeat, reverse repeat, and Dyad repeat) and restriction sites that may interfere with cloning. In addition, to improve translational initiation and performance, Kozak and Shine-Dalgarno Sequences were inserted into the synthetic genes. To increase the efficiency of translational termination, two consecutive stop codons were inserted at the end of cDNAs. The codon usage bias in Human was increased by upgrading the Codon Adaptation Index (CAI) to 0.94. GC content and unfavorable peaks have been optimized to prolong the half-life of the mRNA. The Stem-Loop structures, which impact ribosomal binding and stability of mRNA, were broken. In addition, the optimization process screened and successfully modified those negative cis-acting sites. For the construction of RBD, N/R and S1 constructs, the cDNA corresponding to each region was amplified via PCR by using sequence-specific primers and directionally cloned into the linearized pTK1A-TPA vector by enzymatic restriction PacI/NotI. FL expression vector was generated by In-fusion Cloning System (Takara), amplifying the cDNA by using specific primers overlapping both the synthetic gene and the acceptor empty vector pTK1A. The FL construct was then cloned into the BglII restriction site of pTK1A.

### Transient expression of recombinant SARS-CoV-2 Spike proteins and Western Blotting

HEK293 cells were transiently transfected with SARS-CoV-2-S fragments expression vectors using Lipofectamine 2000 Transfection reagent (Thermo Fisher Scientific). Two days later, the supernatants were collected and concentrated by Amicon Ultra Centrifugal filters (Sigma) and cells were pelleted and lysed in RIPA buffer (Thermo Fisher Scientific). Cell lysates and supernatants were separated by SDS-PAGE and transferred to nitrocellulose membranes. Immunoblotting was performed by using SARS-CoV2 Spike S1 Subunit primary Antibody (Sino Biological) diluted 1:1000 in 5% milk - 0,05% PBS-Tween20. Chemiluminescence detection was performed by using the ECL™ Prime Western Blotting System (Cytiva, Merck) and acquired by ChemiDoc Imaging System (Bio-Rad).

### Recombinant proteins and peptides

RBD-Fc and RBD-6xHis proteins were produced by transient transfection of Expi293F high-density cells with ExpiFectamine 293 Lipid Cation Transfection Reagent (Thermo Fisher) according to the manufacturer’s instructions. The supernatant containing the proteins was collected one week later and subjected to clarification by centrifugation and filtration for the subsequent purification steps. The RBD-Fc protein was purified by affinity chromatography with the AktaPure system with a protein A column (TOYOSCREEN AF-RPROTEIN A HC-650F; Tosoh Bioscience). Briefly, the column was equilibrated with binding buffer (Buffer Phosphate 0.1M pH8) and loaded with the supernatant diluted 1:1 in the same buffer. After washing the column, the protein was recovered by acid elution in 0.1M pH3 citrate buffer, neutralized in Tris-HCl pH9 and subjected to dialysis in PBS1X with slide-A-lyzer (Thermo Fisher) as indicated in the product datasheet. The RBD-6xHIS protein was purified by affinity chromatography of His Tag residues for metals immobilized on the AktaPure system with HisPur ™ Ni-NTA Chromatography Cartridges (Thermo Fisher) column according to the manufacturer’s instructions. Briefly, the column was equilibrated in 5mM PBS1X / Imidazole and loaded with the supernatant diluted 1:1 in the same buffer. After washing, the protein was eluted with PBS1X / Imidazole 0.3M, pH 7.4 and dialyzed in PBS1X with slide-A-lyzers (Thermo Fisher) as indicated in the product datasheet. Once recovered from dialysis, the RBD-Fc and RBD-6xHis proteins were quantified on the spectrophotometer by absorbance at 280nm. Protein purity was evaluated by SDS-PAGE and Western Blot analysis, carried out both in reduced and non-reduced conditions and by standard methods. Lyophilized S protein peptides were purchased from JPT (Berlin, Germany) and resuspended in DMSO at 40 mg/ml. Pools of peptides of 15 aa overlapping by 11 residues were assembled in two pools: pool S1 (residue 1 to 635) and pool S2 (residue 625 to 1273). Peptides and pools were stored at -80°C.

### Production of SARS-CoV2S-pseudoparticles based on VSV

Production of viral pseudoparticles based on vesicular stomatitis virus (VSV) bearing SARS-CoV-2 Spike were produced as previously described^33^. In brief, HEK293T cells were seeded and the next day, transfected with 44 μg plasmid encoding SARS-CoV-2 Spike (pCG1-SARS-2-S, Wuhan Hu-1) using Transit LT-1 (Mirus). The next day, medium was removed, 15 ml fresh medium and then VSV(Fluc_eGFP)-VSV-G added to deliver the defective viral reporter genome (ΔVSV-G, kindly provided by Gert Zimmer, Institute of Virology and Immunology, Mittelhäusern, Switzerland)^34^. After two hours, inoculum was removed, cells washed, and fresh medium added. Pseudoparticles were harvested after 16-24 hours: the supernatant was centrifuged at 1200 rpm 5 min to pellet debris, the supernatant of anti-VSV-G expressing hybridoma cells (Anti-VSV-G antibody (I1, produced from CRL-2700 mouse hybridoma cells, ATCC) added to virus stocks at 1:10 (v/v) to block residual VSV-G containing particles. Supernatants were then concentrated by Vivaspin 20 100 kDa Ultrafiltration devices, immediately aliquoted and frozen at -80°C until use.

### Production of SARS-CoV2S-pseudoparticles based on lentiviral vectors

To generate SARS-CoV-2 lentiviral pseudotype particles, HEK-293TN (System Bioscience) cells were plated in 15-cm dish complete DMEM medium. The following day, 32 μg of reporter plasmid pLenti CMV-GFP-TAV2A-LUC Hygro, 12.5 μg of pMDLg/pRRE (Addgene #12251), 6.25 μg of pRSV-Rev (Addgene #12253), and 9 μg pcDNA3.1_ Spike_del19 were co-transfected following a calcium phosphate transfection. pcDNA3.1_ Spike_del19 (Addgene #155297) was generated by deletion of last 19aa of Spike starting from pcDNA3.1-SARS2-Spike (a gift from Fang Li, Addgene plasmid # 145032). pLenti CMV-GFP-TAV2A-LUC Hygro was generated from pLenti CMV GFP Hygro (Addgene #17446) by addition of T2A-Luciferase by PCR cloning. Twelve hours after transfection, the medium was replaced with complete ISCOVE. 30 h after transfection, the supernatant was collected, clarified by filtration (0.45-μm pore-size) and concentrated by centrifugation for 2h at 20000 rpm. Viral pseudoparticle suspensions were aliquoted and stored at −80°C.

### Animals

BALB/c (H-2^d^) and C57Bl/6 mice (H-2^b^) were purchased from Envigo (Italy). B6.Cg-Tg(K18-ACE2)^2Prlmn/^J mice were purchased from The Jackson Laboratory. Mice were housed under specific pathogen-free conditions and heterozygous mice were used at 6-10 weeks of age. All experimental animal procedures were approved by the Institutional Animal Committee of the San Raffaele Scientific Institute and all infectious work was performed in designed BSL-3 workspaces.

Sixteen, 7-week-old female Sprague-Dawley rats, with a body weight range of 140-155 grams, were purchased from Envigo (Italy).

Sixteen 7-month-old female ferrets (*Mustela putorius furo*) were obtained from a UK Home Office accredited supplier (Highgate Farm, UK). Animals were housed in pairs at Advisory Committee on Dangerous Pathogens (ACDP) containment level 3. Cages met with the UK Home Office *Code of Practice for the Housing and Care of Animals Bred, Supplied or Used for Scientific Procedures* (December 2014). All experimental work was conducted under the authority of a UK Home Office approved project license that had been subject to local ethical review at PHE Porton Down by the Animal Welfare and Ethical Review Body (AWERB) as required by the *Home Office Animals (Scientific Procedures) Act 1986*.

### Vaccination

#### Mice

DNA-EGT was performed in mice quadriceps injected with doses ranging from 0.1 µg to 20 µg and electrically stimulated as previously described ^10^. The DNA was formulated in Phosphate Buffered Saline (PBS) at a concentration of 0.2 mg/ml. DNA-EP was performed with an IGEA Cliniporator (Carpi, Italy), using a needle electrode (electrode A-15-4B). At different time points, antibody and cell mediated immune response were analyzed.

#### Rats

After a suitable quarantine period, animals were divided in three different experimental groups (4 females/group) and immunized by intramuscular electroporation, alternating quadriceps at each vaccine administration.

#### Ferrets

Eight ferrets were immunized twice at day -42 and -14 with 400µg RBD intra-muscularly in the quadriceps muscle of the right leg using followed by electroporation. An additional eight ferrets remained unvaccinated.

### Viruses and *in vivo* treatments

The hCoV-19/Italy/LOM-UniSR-1/2020 (GISAID Accession ID: EPI_ISL_413489) isolate of SARS-CoV-2 was used in this study. Virus isolation studies were carried out in BSL-3 workspace and performed in Vero E6 cells, which were cultured at 37°C, 5% CO_2_ in complete medium (DMEM supplemented with 10% FBS, 1% penicillin plus streptomycin, 1% L-glutamine). Virus stocks were titrated using both Plaque Reduction Assayurn (PRA, PFU/ml) and Endpoint Dilutions Assay (EDA, TCID_50_/ml). In PRA, confluent monolayers of Vero E6 cells were infected with eight 10-fold dilutions of virus stock. After 1 h of adsorption at 37°C, the cell-free virus was removed. Cells were then incubated for 48 h in DMEM containing 2% FBS and 0.5% agarose. Cells were fixed and stained, and viral plaques were counted. In EDA, Vero E6 cells were seeded into 96 wells plates and infected at 95% of confluency with base 10 dilutions of virus stock. After 1 h of adsorption at 37°C, the cell-free virus was removed, cells were washed with PBS 1X, and complete medium was added to cells. After 48 h, cells were observed to evaluate the presence of a cytopathic effect (CPE). TCID_50_/ml of viral stocks were then determined by applying the Reed–Muench formula.

K18-hACE2 mice were immunized with 10 ug of COVID-*e*Vax or saline solution twice 21 days apart intra-muscularly followed by electroporation as described above.

Virus infection was performed via intranasal administration of 1 x 10^5^ TCID_50_ per mouse under Isoflurane 2% (# IsoVet250) anesthesia. Mice were monitored to record body weight, clinical and respiratory parameters.

Eight ferrets were immunized twice at day -42 and -14 with 400µg RBD intra-muscularly in the quadriceps muscle of the right leg using followed by electroporation as described above. An additional eight ferrets remained unvaccinated. All ferrets were challenged with SARS-CoV-2 Victoria/01/2020^35^ six weeks following first vaccination. Challenge virus (5×10^6^ PFU/ml) was delivered by intranasal instillation (1.0 ml total, 0.5 ml per nostril) diluted in phosphate buffered saline (PBS). Nasal washes and throat swabs were taken from all ferrets at 2, 4, 6- and 7-days post challenge. Nasal washes were obtained by flushing the nasal cavity with 2 ml PBS. For throat swabs, a flocked swab (MWE Medical Wire, Corsham, UK) was gently stroked across the back of the pharynx in the tonsillar area. Four animals from each group were euthanized at day 3 and the remaining animals from each group were euthanized at day 7.

### Luciferase assay

BALB/c mice (5 mice/group) were anesthetized using 97% oxygen and 3% isoflurane (Isoba, MSD animal Health, Walton, UK) then injected by DNA electroporation with a DNA plasmid encoding Luciferase (pcDNA3-Hygro-Luc 1 μg/mouse) expressing luciferase in a 50 μl volume in quadriceps muscle. Mice were electroporated by means of a Cliniporator Device EPS01 N-10-4B electrodes with the following electrical conditions in Electro-Gene-Transfer (EGT) modality: 8 pulses 20 msec each at 110V, 8Hz, 120msec interval. Imaging was performed under gas anesthesia at Xenogen IVIS 200 at 48h after injection, 8 min after injecting s.c. a luciferin solution (15mg/ml, Perkin Elmer) at 10 μl/g of body weight.

### ELISpot assays

For the B cell Elispot assay, pools of sera collected from mice vaccinated with RBD or FL constructs were tested against each of the 338 peptides covering the entire Spike protein, pre-coated on 96 well plate, in order to identify the linear epitopes. Sequences of positive hits were then mapped on three-dimensional structure of Spike protein, hence outlining the epitope domains. The T cell ELISPOT for mouse IFNγ was performed as previously described^36^. RBD peptides are 132 out of the 338 peptides covering the whole Spike protein (from peptide nr.4 to peptide nr.136). In order to identify immunodominant RBD epitopes (here highlighted in yellow), Elispot assay was performed by stimulating splenocytes from RBD vaccinated Balb/c mice for 20h with RBD peptide pools. Pools (from 1 to 24) were distributed as a matrix (the intersection of two pools identifies one RBD peptide), with each pool comprising up to 12 RBD peptides. Immunodominant RBD peptides were identified at the intersection of pools showing >50 SFCs.

### Antibody detection assays

Antibody titration was performed both on sera, obtained by retro-orbital bleeding, and on bronchoalveolar lavages (BALs), obtained by flushing 1ml PBS in the lungs. The ELISA plates were functionalized by coating with the RBD-6xHis protein at a concentration of 1 μg/ml and incubated about 18 hours at 4°C. Subsequently the plates were blocked with 3% BSA / 0.05% Tween-20 / PBS for 1 hour at room temperature and then the excess solution was eliminated. The sera of the immunized mice were then added at a dilution of 1/300 and diluted 1:3 up to 1/218,700, in duplicate, and the plates incubated for 2 hours at room temperature. After a double wash with 0.05% Tween-20 / PBS, the secondary anti-murine IgG or anti-murine IgM conjugated with alkaline phosphatase was added and the plates were incubated for 1 hour at room temperature. After a double wash with 0.05% Tween-20 / PBS, the binding of the secondary was detected by adding the substrate for alkaline phosphatase and measuring the absorbance at 405nm by means of an ELISA reader after incubation for 2 hours. IgG antibody titers against the S protein RBD were evaluated at several time points. Regarding antibody titration on sera of vaccinated K18-hACE2 mice, the sera were added at a dilution of 1/300 to 1/218700, in duplicate, and the plates incubated O/N at 4°C. After three washes with 0.05% Tween-20 / PBS, the secondary anti-murine IgG HRP (1:2000) was added and the plates were incubated for 1 hour at room temperature. After a wash with 0.05% Tween-20 / PBS, the binding of the secondary was detected by adding TMB substrate reagent (Biosciences). The reaction was blocked with 0,5M H_2_SO_4_ and the absorbance at 450 nm and reference 630 nm was measured.

For neutralization experiments with SARS-CoV-2 VSV-based pseudoparticles, Caco-2 cells were seeded in 96-well plates in 180 μl medium at 10,000 cells/well. The next day, serially diluted heat-inactivated sera in PBS or diluted BALs were added to CoV2S-PPs at 1:1 dilution with max. 10% serum or BAL on PPs. Sera/BAL-PP mixtures were incubated 1h at 37°C, then added to Caco-2 cells in duplicates at 1:10 dilution (max. 1% serum on cells) and cells incubated at 37°C. 16h post-transduction, Firefly luciferase activity was measured using the Promega Luciferase Assay System (E1501), values normalized to PPs treated with PBS only. IC_50_ fit by Prism Inhibitor vs. Normalized response (variable slope); samples with < 50% inhibition at 10% serum were excluded. These assays have been performed in a BSL-2 facility at Ulm University.

For lentiviral pseudotype neutralization assay, HEK293TN-hACE2 were plated at 10^4^ cells/well in white 96-well plates (100 μl / well of complete DMEM medium). The next day, cells were infected with 0.1 MOI of SARS-CoV-2-GFP/luciferase pseudoparticles that were subjected to preincubation with serially diluted sera. In detail, sera samples were serially diluted three-fold in PBS in order to obtain a 7-point dose-response curve (plus PBS alone as untreated control). Thereafter, aliquots of undiluted or three-fold serially-diluted sera were further diluted 1:10 in aliquots of SARS-CoV-2 pseudoparticles adjusted to contain 0.1 M.O.I. / 50 μl of complete culture medium. After incubation for 1h at 37°C, 50 μl of serum/SARS-CoV-2 pseudoparticles mixture was added to each well and plates were incubated for 24h at 37°C. Thus, the starting serum dilutions was 1:30. Each dilution was tested in triplicate. After 24h of incubation cell infection was measured by luciferase assay using Bright-Glo™ Luciferase System (Promega) Infinite F200 plate reader (Tecan) was used to read luminescence. Obtained RLUs were normalized to controls and dose response curves were generated by nonlinear regression curve fitting with GraphPad Prism to calculate Neutralization Dose 50 (ND_50_).

HEK293TN-hACE2 were generated by transduction of HEK293TN with a lentiviral vector engineered to stably express hACE2 (described elsewhere, paper submitted).

To test the ability of elicited antibodies to neutralize the virus *in vitro*, Vero E6 cells (20,000 cells / well) were seeded 24 hours prior to infection in 96 well plates (Costar). Serum samples from mice were incubated at 56°C for 30 minutes and then serially two-fold diluted in cell culture medium (10- to 10240-fold). Serum dilutions were then mixed to 100 TCID_50_ of SARS-CoV-2 (virus isolated in January 2020 at the National Institute for Infectious Diseases “L. Spallanzani” in Rome, 2019-nCoV/Italy-INMI1, clade V strain; GISAID accession number: EPI_ISL_410545) in 96-well plates and incubated at 37°C for 30 minutes in 5% vol / vol CO2. The virus/serum mixture was then added to the cells and incubated at 37°C for microscopic examination of the cytopathic effect after 48 hours. Cell supernatants were then removed, and cells were fixed/stained for 30 minutes with a solution of 2% formaldehyde (AppliChem, Darmstadt, Germany) in Crystal Violet (Diapath S.p.A., Martinengo, Bergamo, Italy). The fixing solution was removed, and cell viability measured by photometer at 595 nm (Synergy HTX; BioTek Instruments, Winooski, VT, USA). The serum dilution inhibiting 50% of the CPE (IC50) was calculated using Prism 7 (GraphPad Software, San Diego, CA, USA) as described ^37^. Tests were performed in duplicate with negative control samples from unvaccinated mice and positive control samples from a COVID-19 patient with known neutralizing titer. The same assay was repeated with the SARS-CoV-2 strain belonging to the G clade (SARS-CoV-2/Human/ITA/PAVIA10734/2020, clade G, D614G (S), in GISAID EPI_ISL_568579; isolated at Policlinico San Matteo in the Laboratory of Prof. Fausto Baldanti). These assays have been performed in a BSL-3 facility at the Spallanzani Institute in Rome.

### Cell Isolation and Flow Cytometry

In experiments performed with vaccinated WT mice, the intracellular staining was performed according to the procedure described in Giannetti et al^38^. Briefly, PBMC or splenocytes were treated with ACK Lysing buffer (Life Technologies) for red blood cell lysis and resuspended in 0.6ml RPMI, 10% FCS and incubated with the indicated pool of peptides (5 μg/ml final concentration of each peptide) and brefeldin A (1 μg/ml; BD Pharmingen) at 37°C for 12-16 hours. Cells were then washed and stained with surface antibodies. After washing, cells were fixed, permeabilized and incubated with anti-IFNγ (XMG1.2) and -TNF*α* (MP6-XT22; all from eBioscience, USA), fixed with 1% formaldehyde in PBS and analyzed on a CytoFLEX flow cytometer. DMSO and PMA/IONO (Sigma) at 10μg/ml were used as internal negative and positive control of the assay, respectively.

In experiments performed with SARS-CoV-2-infected mice, lung was perfused through the right ventricle with PBS at the time of autopsy and after the brain was removed from the skull. Lung tissue was digested in RPMI 1640 containing 3.2 mg/ml Collagenase IV (Sigma) and 25 U/ml DNAse I (Sigma) for 30 minutes at 37°C. Brain was digested in RPMI 1640 containing 1 mg/ml Collagenase D (Sigma) and 6,3 μg/ml DNAse I (Sigma) for 30 minutes at 37°C. Homogenized lung and brain were passed through 70 μm nylon mesh to obtain a single cell suspension. Cells were resuspended in 36% percoll solution (Sigma) and centrifuged for 20 minutes at 2000 rpm (light acceleration and low brake). The remaining red blood cells were removed with ACK lysis. For analysis of *ex-vivo* intracellular cytokine production, 1 mg/ml of brefeldin A (Sigma) was included in the digestion buffer. All flow cytometry stainings of surface-expressed and intracellular molecules were performed as described^39^. Briefly, cells were stimulated for 4 h at 37°C with 15-mer peptides overlapping by 11 aminoacids (5 μg/ml) covering the receptor-biding-domain (RBD) of SARS-CoV-2. Cell viability was assessed by staining with Viobility™ 405/520 fixable dye (Miltenyi). Antibodies (Abs) used are indicated in the table below.

| Name | Clone | Source and catalog number |
| --- | --- | --- |
| CD8α | 53-6.7 | Biolegend 100723 |
| CD3 | 145-2C11 | Pharmingen 552774 |
| CD44 | IM7 | Biolegend 103028 |
| IFNγ | XMG1.2 | Biolegend 505830 |
| TNFα | MP6-XT22 | Biolegend 506329 |
| CD62L | MEL-14 | Biolegend 104453 |
| CD4 | RM4-5 | BD Biosciences 740208 |
| IL-17A | TC11-18H10 | BD Pharmingen 562542 |
| IL-5 | TRFK5 | BD 554395 |
| IL-4 | 11B11 | BD Pharmingen 554436 |

Flow cytometry analysis was performed on CytoFLEX LX Beckton Coulter and analyzed with FlowJo software (Treestar).

### Tissue homogenates and viral titers

Tissues homogenates were prepared by homogenizing perfused lung using gentleMACS Octo Dissociator (Miltenyi) in M tubes containing 1 ml of DMEM 0%. Samples were homogenized for three times with the program m_Lung_01_02 (34 seconds, 164 rpm). The homogenates were centrifuged at 3500 rpm for 5 minutes at 4°C. The supernatant was collected and stored at −80°C for viral isolation and viral load detection. Viral titer was calculated by 50% tissue culture infectious dose (TCID_50_). Briefly, Vero E6 cells were seeded at a density of 1.5 × 10^4^ cells per well in flat-bottom 96-well tissue culture plates. The following day, 2-fold dilutions of the homogenized tissue were applied to confluent cells and incubated 1 h at 37°C. Then, cells were washed with phosphate-buffered saline (PBS) and incubated for 72 h at 37°C in DMEM 2% FBS. Cells were fixed with 4% paraformaldehyde for 20 min and stained with 0.05% (wt/vol) crystal violet in 20% methanol.

### RNA extraction and qPCR

Tissues homogenates were prepared by homogenizing perfused lung using gentleMACS dissociator (Miltenyi) with the RNA_02 program in M tubes in 1 ml of Trizol (Invitrogen). The homogenates were centrifuged at 2000 g for 1 min at 4°C and the supernatant was collected. RNA extraction was performed by combining phenol/guanidine-based lysis with silica membrane-based purification. Briefly, 100 μl of Chloroform was added to 500 μl of homogenized sample; after centrifugation, the aqueous phase was added to 1 volume of 70% ethanol and loaded on ReliaPrep™ RNA Tissue Miniprep column (Promega, Cat #Z6111). Total RNA was isolated according to the manufacturer’s instructions. qPCR was performed using TaqMan Fast virus 1 Step PCR Master Mix (Life Technologies), standard curve was drawn with 2019_nCOV_N Positive control (IDT), the primers used were: 2019-nCoV_N1-Forward Primer (5’-GAC CCC AAA ATC AGC GAA AT-3’), 2019-nCoV_N1-Reverse Primer (5’-TCT GGT TAC TGC CAG TTG AAT CTG-3’) 2019-nCoV_N1-Probe (5’-FAM-ACC CCG CAT TAC GTT TGG ACC-BHQ1-3’) (Centers for Disease Control and Prevention (CDC) Atlanta, GA 30333). All experiments were performed in duplicate.

### Whole-body plethysmography

Whole-body plethysmography (WBP) was performed using WBP chamber (DSI Buxco respiratory solutions, DSI). First mice were allowed to acclimate inside the chamber for 10 minutes, then respiratory parameters were acquired for 15 minutes using FinePointe software.

### Confocal immunofluorescence histology and histochemistry and N-SARS-CoV-2 signal quantification

Lungs of infected mice were collected and fixed in 4% paraformaldehyde (PFA). Samples were then dehydrated in 30% sucrose prior to embedding in OCT freezing media (Bio-Optica). Twenty micrometer sections were cut on a CM1520 cryostat (Leica) and adhered to Superfrost Plus slides (Thermo Scientific). Sections were then permeabilized and blocked in PBS containing 0.3% Triton X-100 (Sigma-Aldrich) and 5% FBS followed by staining in PBS containing 0.3% Triton X-100 and 1% FBS. Slides were stained for SARS-CoV-2 nucleocapsid (GeneTex) for 1h RT. Then, slides were stained with Alexa Fluor 568 Goat Anti-Rabbit antibody for 2h RT. All slides were analyzed by confocal fluorescence microscopy (Leica TCS SP5 Laser Scanning Confocal). For SARS-CoV-2 N protein immunohistochemistry, mice were perfused with PBS and brains were collected in Zn-formalin and transferred into 70% ethanol 24 h later. Tissue was then processed, embedded in paraffin and automatically stained for SARS-CoV-2 (2019-nCoV) Nucleocapsid Antibody (SINO BIO, 40143-R019) through LEICA BOND RX 1h RT and developed with Bond Polymer Refine Detection (Leica, DS9800). Brightfield images were acquired through an Aperio Scanscope System CS2 microscope and an ImageScope program (Leica Biosystem) following the manufacturer’s instructions. In both immunofluorescence and histochemistry, SARS-CoV-2 N protein percentage of positive area was determined by QuPath (Quantitative Pathology & Bioimage Analysis) software.

### Statistical analyses and software

Detailed information concerning the statistical methods used is provided in the figure legends. Flow cytometry data were collected using FlowJo Version 10.5.3 (Treestar). Statistical analyses were performed with GraphPad Prism software version 8 (GraphPad). Immunofluorescence and histochemical imaging quantifications were performed with Aperio Scanscope System and QuPath software (Quantitative Pathology & Bioimage Analysis). *n* represents individual mice analyzed per experiments. Error bars indicate the standard error of the mean (SEM). We used Mann-Whitney U-tests to compare two groups with non-normally distributed continuous variables. We used two-way ANOVA followed by Sidak’s multiple comparisons tests to analyze experiments with multiple groups and two independent variables. Significance is indicated as follows: *p<0.05; **p<0.01. Comparisons are not statistically significant unless indicated.

## Supplementary Figure Legends

**Supplementary Figure 1.**
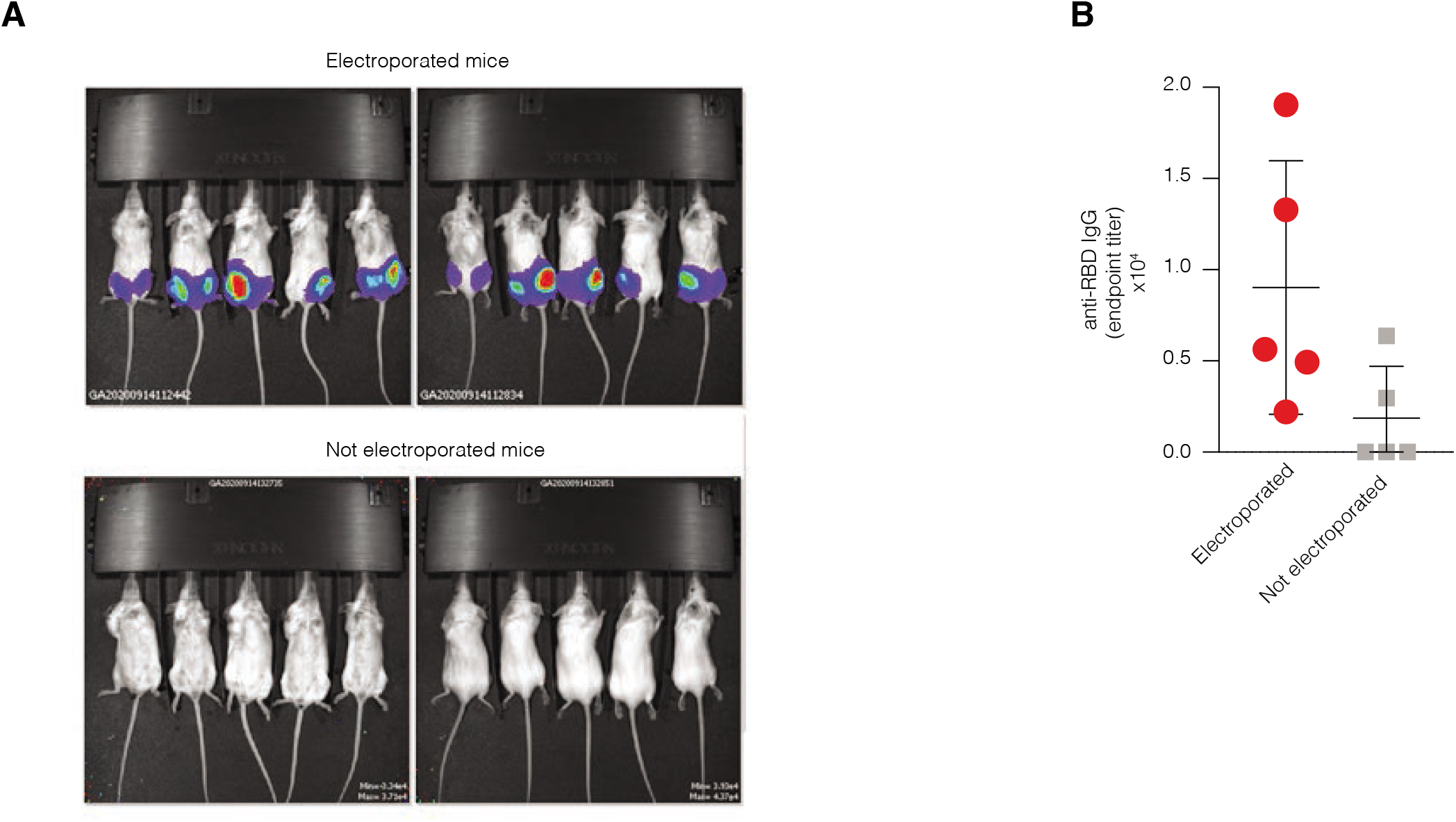
Electroporation increases the level of gene expression upon DNA immunization. **(A)** BALB/c mice (*n* = 5) were either injected i.m. with 1 mg of a plasmid expressing firefly luciferase followed by electroporation (upper panel, electroporated mice) or not (lower panel, non-electroporated mice). Forty-eight hours later, optical imaging was carried out using an IVIS 200 system. Ventral and dorsal images were taken. **(B)** BALB/c mice were injected with 5 mg of RBD vaccine, with or without electroporation. 14 days later mice were bled, and anti-RBD IgG endpoint titers were measured by ELISA.

**Supplementary Figure 2.**
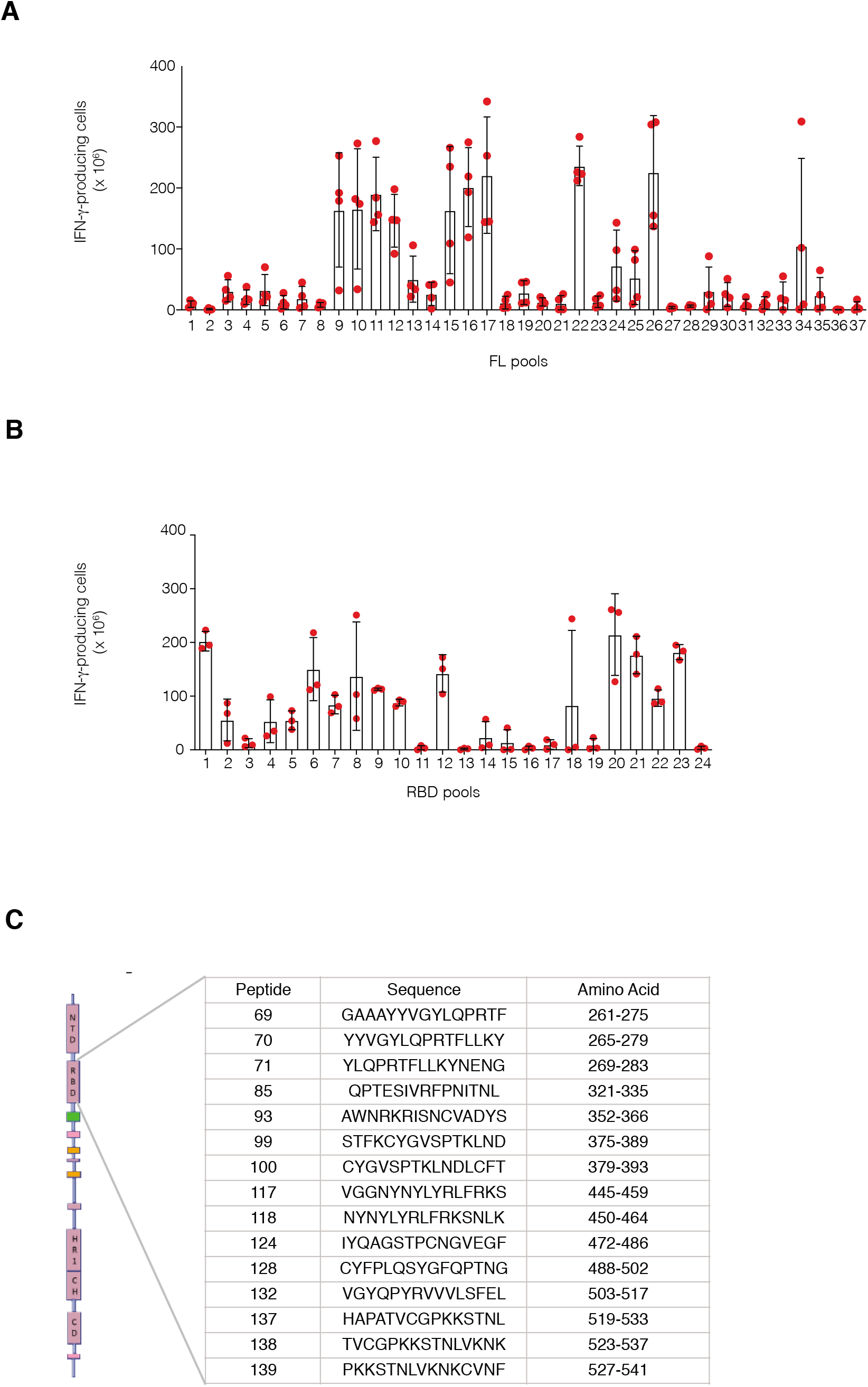
T cell epitope mapping in RBD vaccinated mice. **(A-B)** IFNγ^+^ T cell response measured by ELISpot assay on splenocytes collected from FL or RBD-vaccinated BALB/c mice, following stimulation with matrix mapping FL or RBD peptide pools. **(C)** Schematic representation of the SARS-CoV-2 Spike protein and identification of immunodominant peptides in BALB/c mice.

**Supplementary Figure 3.**
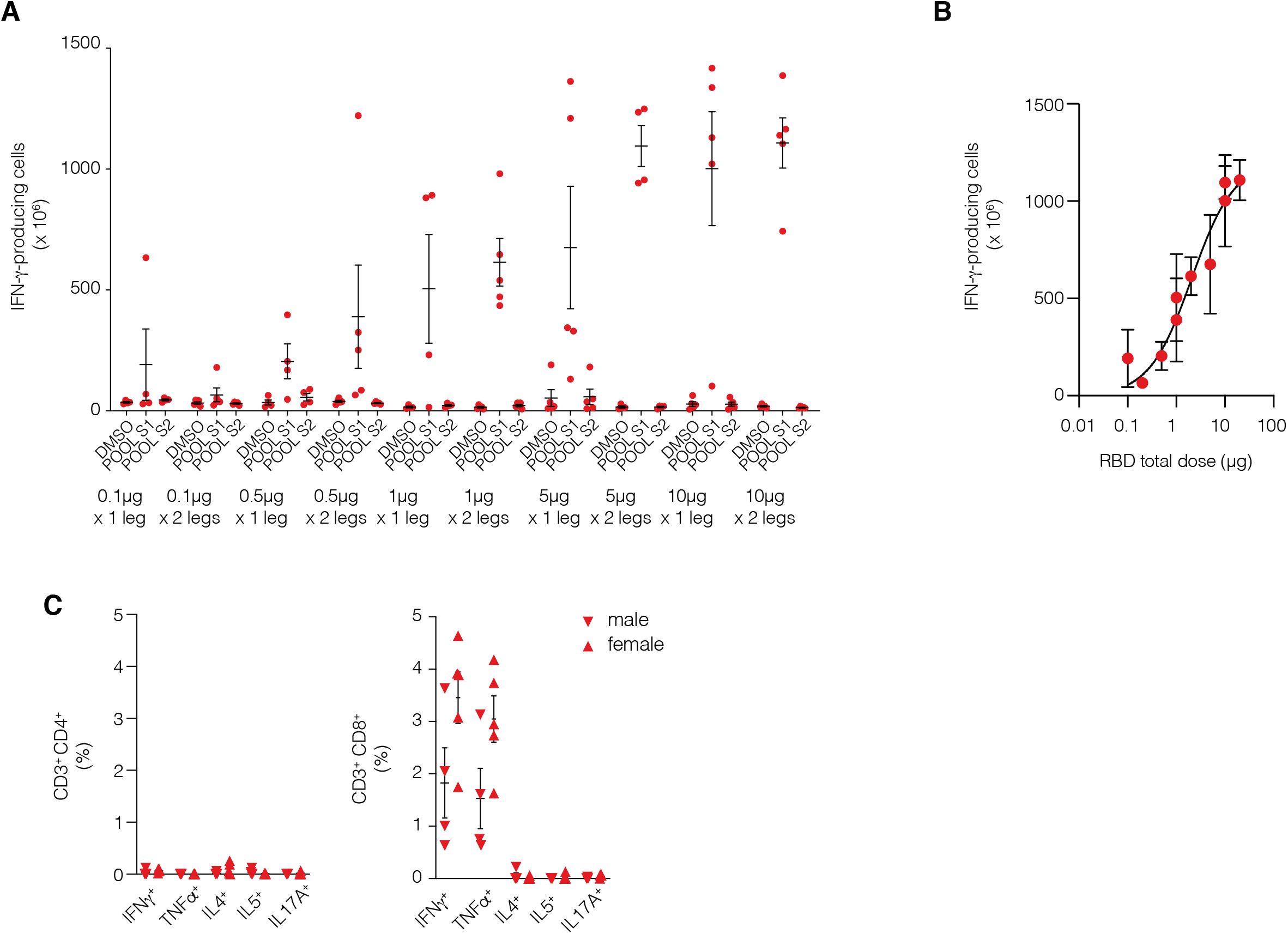
RBD-specific immune response in RBD vaccinated C57BL/6 mouse model. **(A)** IFN-γ^+^ T cell response measured by ELISpot assay on splenocytes collected at day 38 from C57BL/6 mice vaccinated with increasing doses of RBD vaccine (from 0.1 to 20 μg, administered at one or two sites) and restimulated with Spike peptide pools S1 and S2. **(B)** Non-linear fitting curve of the dose-response against RBD pool S1 peptides, measured by means of ELISpot assay performed on splenocytes from RBD-vaccinated C57BL/6 mice. **(C)** T cell characterization by intracellular staining on PBMCs collected from males and females vaccinated mice (administered dose: 5μg / leg).

**Supplementary Figure 4.**
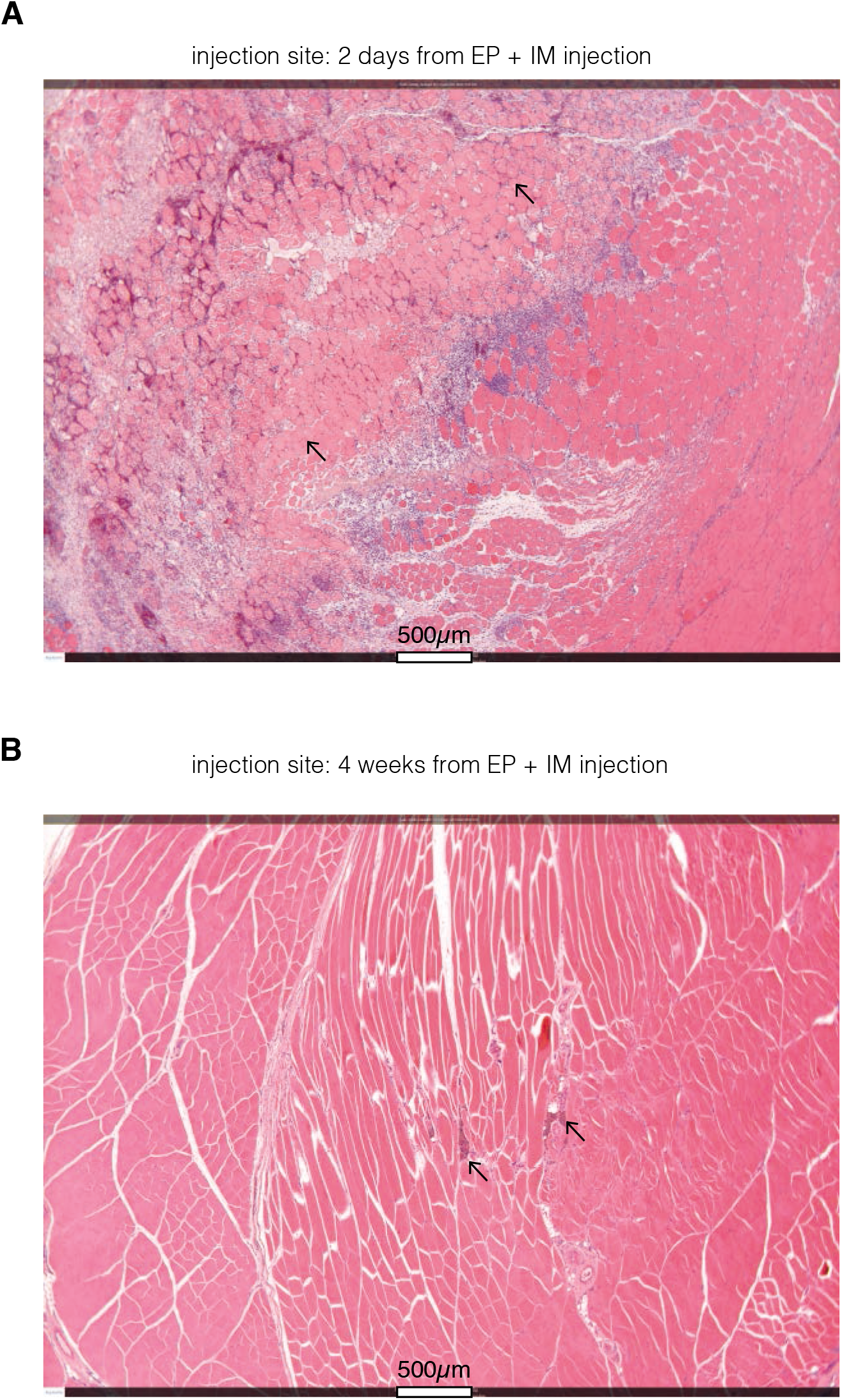
Histopathological evaluation of electroporated tissues in rat model. **(A)** Histological section of the left injection site in a 400 µg RBD-vaccinated rat performed two days after the third and last DNA injection (i.e. day 30). Arrows indicate the necrosis of muscle fibers, surrounded by inflammatory reaction (i.e., polymorphonuclear cells, mixed mononuclear cell infiltration, predominantly macrophages). The cavities surrounded by the necrotic carbonized muscle fibers are suggested to be related to the electroporation procedure. The lesions were mostly scored as mild to moderate and were similar in all the groups. **(B)** Histological section of the left injection site in a 400 µg RBD vaccinated rat performed 4 weeks after third and last DNA injection (i.e. day 57). Arrows indicate brownish pigmented muscle fibers, probably related to a minimal chronic inflammation due to the electroporation procedure. This image demonstrates a complete recovery of the injection site lesions at this stage, in comparison to (A).

**Supplementary Figure 5.**
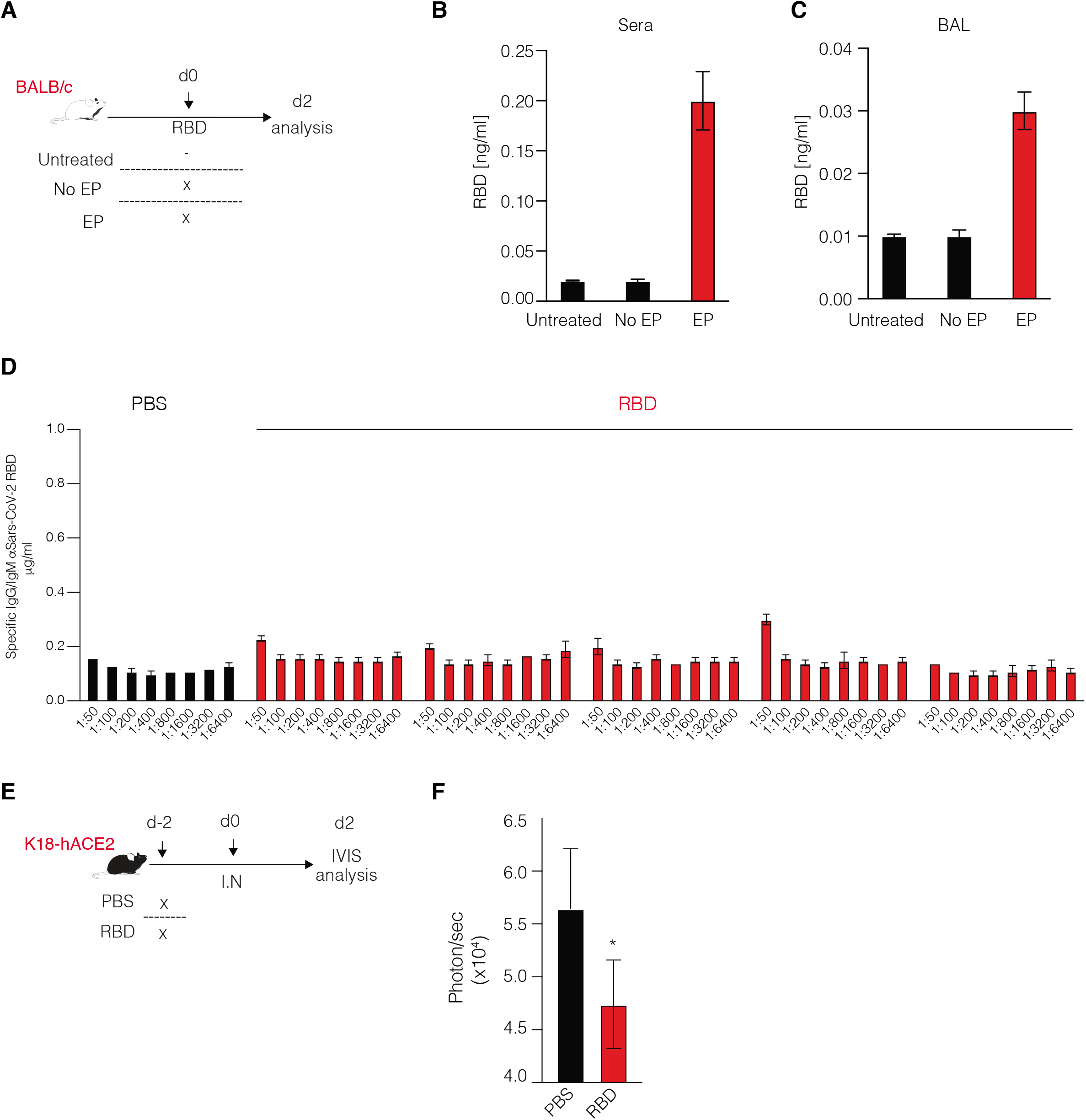
Assessment of secreted RBD in RBD vaccinated mice. **(A)** Schematic representation of the experimental setup. BALB/c mice were vaccinated with 20 μg of RBD vaccine, with or without electroporation, and 48 hours later the secretion of RBD protein was assessed in sera and BALs. **(B)** Measurement of secreted RBD protein in sera from control mice and RBD vaccinated mice, with or without EP. **(C)** Measurement of secreted RBD protein in BALs from same groups of mice as in (B). **(D)** Measurement of anti-RBD antibodies in the sera at day 2 after RBD or PBS vaccination **(E)** Schematic representation of the experimental setup. K18-hACE2 mice were vaccinated with 20 μg of RBD vaccine and 2 days later a lentiviral vector pseudotyped with the SARS-CoV-2 spike protein and encoding for luciferase RBD protein was intranasally administered. Two days later the lungs of treated mice were assessed for bioluminescence using an in vivo imaging system. **(F)** Comparison of bioluminescence assessed by means of *in vivo* imaging system in control K18-hACE2 mice and K18-hACE2 RBD vaccinated mice. * p value < 0.05

## Supplementary Table Legends

**Table S1.***List of immunodominant B epitopes.*

**Table S2.***Scheme of RBD peptide pool matrix.*

## References

1. Krammer, F. (2020). SARS-CoV-2 vaccines in development. Nature 586, 516–527.

2. Gebre, M.S., Brito, L.A., Tostanoski, L.H., Edwards, D.K., Carfi, A., and Barouch, D.H. (2021). Novel approaches for vaccine development. Cell 184, 1589–1603.

3. Conforti, A., Marra, E., Roscilli, G., Palombo, F., Ciliberto, G., and Aurisicchio, L. (2020). Are Genetic Vaccines the Right Weapon against COVID-19? Mol Ther 28, 1555–1556.

4. Yang, Z., Kong, W., Huang, Y., Roberts, A., Murphy, B.R., Subbarao, K., and Nabel, G.J. (2004). A DNA vaccine induces SARS coronavirus neutralization and protective immunity in mice. Nature 428, 561–564.

5. Modjarrad, K., Roberts, C.C., Mills, K.T., Castellano, A.R., Paolino, K., Muthumani, K., Reuschel, E.L., Robb, M.L., Racine, T., Oh, M., et al. (2019). Safety and immunogenicity of an anti-Middle East respiratory syndrome coronavirus DNA vaccine: a phase 1, open-label, single-arm, dose-escalation trial. Lancet Infect Dis 19, 1013–1022.

6. Wrapp, D., Wang, N., Corbett, K.S., Goldsmith, J.A., Hsieh, C.-L., Abiona, O., Graham, B.S., and McLellan, J.S. (2020). Cryo-EM structure of the 2019-nCoV spike in the prefusion conformation. Science 367, 1260–1263.

7. Wang, L., Shi, W., Chappell, J.D., Joyce, M.G., Zhang, Y., Kanekiyo, M., Becker, M.M., Doremalen, N. van, Fischer, R., Wang, N., et al. (2018). Importance of Neutralizing Monoclonal Antibodies Targeting Multiple Antigenic Sites on the Middle East Respiratory Syndrome Coronavirus Spike Glycoprotein To Avoid Neutralization Escape. J Virol 92, e02002–17.

8. Piccoli, L., Park, Y.-J., Tortorici, M.A., Czudnochowski, N., Walls, A.C., Beltramello, M., Silacci-Fregni, C., Pinto, D., Rosen, L.E., Bowen, J.E., et al. (2020). Mapping Neutralizing and Immunodominant Sites on the SARS-CoV-2 Spike Receptor-Binding Domain by Structure-Guided High-Resolution Serology. Cell 183, 1024–1042.e21.

9. Fattori, E., Monica, N.L., Ciliberto, G., and Toniatti, C. (2002). Electro-Gene-Transfer: A New Approach for Muscle Gene Delivery. Somat Cell Molec Gen 27, 75–83.

10. Cappelletti, M., Zampaglione, I., Rizzuto, G., Ciliberto, G., Monica, N.L., and Fattori, E. (2003). Gene electro-transfer improves transduction by modifying the fate of intramuscular DNA. J Gene Medicine 5, 324–332.

11. Aurisicchio, L., Mancini, R., and Ciliberto, G. (2014). Cancer vaccination by electro-gene-transfer. Expert Rev Vaccines 12, 1127–1137.

12. Hojman, P. (2010). Basic Principles and Clinical Advancements of Muscle Electrotransfer. Curr Gene Ther 10, 128–138.

13. Doremalen, N. van, Lambe, T., Spencer, A., Belij-Rammerstorfer, S., Purushotham, J.N., Port, J.R., Avanzato, V.A., Bushmaker, T., Flaxman, A., Ulaszewska, M., et al. (2020). ChAdOx1 nCoV-19 vaccine prevents SARS-CoV-2 pneumonia in rhesus macaques. Nature 586, 578–582.

14. Huo, J., Bas, A.L., Ruza, R.R., Duyvesteyn, H.M.E., Mikolajek, H., Malinauskas, T., Tan, T.K., Rijal, P., Dumoux, M., Ward, P.N., et al. (2020). Neutralizing nanobodies bind SARS-CoV-2 spike RBD and block interaction with ACE2. Nat Struct Mol Biol 27, 846–854.

15. Schafer, K.A., Eighmy, J., Fikes, J.D., Halpern, W.G., Hukkanen, R.R., Long, G.G., Meseck, E.K., Patrick, D.J., Thibodeau, M.S., Wood, C.E., et al. (2018). Use of Severity Grades to Characterize Histopathologic Changes. Toxicol Pathol 46, 256–265.

16. McCray, P.B., Pewe, L., Wohlford-Lenane, C., Hickey, M., Manzel, L., Shi, L., Netland, J., Jia, H.P., Halabi, C., Sigmund, C.D., et al. (2007). Lethal Infection of K18-hACE2 Mice Infected with Severe Acute Respiratory Syndrome Coronavirus▿. J Virol 81, 813–821.

17. Zheng, J., Wong, L.-Y.R., Li, K., Verma, A.K., Ortiz, M.E., Wohlford-Lenane, C., Leidinger, M.R., Knudson, C.M., Meyerholz, D.K., McCray, P.B., et al. (2021). COVID-19 treatments and pathogenesis including anosmia in K18-hACE2 mice. Nature 589, 603–607.

18. Menachery, V.D., Gralinski, L.E., Baric, R.S., and Ferris, M.T. (2015). New Metrics for Evaluating Viral Respiratory Pathogenesis. Plos One 10, e0131451.

19. Dinnon, K.H., Leist, S.R., Schäfer, A., Edwards, C.E., Martinez, D.R., Montgomery, S.A., West, A., Yount, B.L., Hou, Y.J., Adams, L.E., et al. (2020). A mouse-adapted model of SARS-CoV-2 to test COVID-19 countermeasures. Nature 586, 560–566.

20. Jorritsma, S.H.T., Gowans, E.J., Grubor-Bauk, B., and Wijesundara, D.K. (2016). Delivery methods to increase cellular uptake and immunogenicity of DNA vaccines. Vaccine 34, 5488– 5494.

21. Diaz, C.M., Chiappori, A., Aurisicchio, L., Bagchi, A., Clark, J., Dubey, S., Fridman, A., Fabregas, J.C., Marshall, J., Scarselli, E., et al. (2013). Phase 1 studies of the safety and immunogenicity of electroporated HER2/CEA DNA vaccine followed by adenoviral boost immunization in patients with solid tumors. J Transl Med 11, 62.

22. Aurisicchio, L., Fridman, A., Mauro, D., Sheloditna, R., Chiappori, A., Bagchi, A., and Ciliberto, G. (2020). Safety, tolerability and immunogenicity of V934/V935 hTERT vaccination in cancer patients with selected solid tumors: a phase I study. J Transl Med 18, 39.

23. Peruzzi, D., Gavazza, A., Mesiti, G., Lubas, G., Scarselli, E., Conforti, A., Bendtsen, C., Ciliberto, G., Monica, N.L., and Aurisicchio, L. (2010). A Vaccine Targeting Telomerase Enhances Survival of Dogs Affected by B-cell Lymphoma. Mol Ther 18, 1559–1567.

24. Impellizeri, J.A., Gavazza, A., Greissworth, E., Crispo, A., Montella, M., Ciliberto, G., Lubas, G., and Aurisicchio, L. (2018). Tel-eVax: a genetic vaccine targeting telomerase for treatment of canine lymphoma. J Transl Med 16, 349.

25. Aurisicchio, L., Salvatori, E., Lione, L., Bandini, S., Pallocca, M., Maggio, R., Fanciulli, M., Nicola, F.D., Goeman, F., Ciliberto, G., et al. (2019). Poly-specific neoantigen-targeted cancer vaccines delay patient derived tumor growth. J Exp Clin Canc Res 38, 78.

26. Impellizeri, J.A., Ciliberto, G., and Aurisicchio, L. (2014). Electro-gene-transfer as a new tool for cancer immunotherapy in animals. Vet Comp Oncol 12, 310–318.

27. Gary, E.N., and Weiner, D.B. (2020). DNA vaccines: prime time is now. Curr Opin Immunol 65, 21–27.

28. Martin, J.E., Louder, M.K., Holman, L.A., Gordon, I.J., Enama, M.E., Larkin, B.D., Andrews, C.A., Vogel, L., Koup, R.A., Roederer, M., et al. (2008). A SARS DNA vaccine induces neutralizing antibody and cellular immune responses in healthy adults in a Phase I clinical trial. Vaccine 26, 6338–6343.

29. Smith, T.R.F., Patel, A., Ramos, S., Elwood, D., Zhu, X., Yan, J., Gary, E.N., Walker, S.N., Schultheis, K., Purwar, M., et al. (2020). Immunogenicity of a DNA vaccine candidate for COVID-19. Nat Commun 11, 2601.

30. Greaney, A.J., Loes, A.N., Crawford, K.H.D., Starr, T.N., Malone, K.D., Chu, H.Y., and Bloom, J.D. (2021). Comprehensive mapping of mutations in the SARS-CoV-2 receptor-binding domain that affect recognition by polyclonal human plasma antibodies. Cell Host Microbe 29, 463–476.e6.

31. Tortorici, M.A., Czudnochowski, N., Starr, T.N., Marzi, R., Walls, A.C., Zatta, F., Bowen, J.E., Jaconi, S., Iulio, J.D., Wang, Z., et al. (2021). Broad sarbecovirus neutralization by a human monoclonal antibody. Nature, 1–6.

32. Jarjour, N.N., Masopust, D., and Jameson, S.C. (2020). T cell memory: Understanding COVID-19. Immunity 54, 14–18.

33. Hoffmann, M., Arora, P., Groß, R., Seidel, A., Hörnich, B.F., Hahn, A.S., Krüger, N., Graichen, L., Hofmann-Winkler, H., Kempf, A., et al. (2021). SARS-CoV-2 variants B.1.351 and P.1 escape from neutralizing antibodies. Cell 184, 2384–2393.e12.

34. Rentsch, M.B., and Zimmer, G. (2011). A Vesicular Stomatitis Virus Replicon-Based Bioassay for the Rapid and Sensitive Determination of Multi-Species Type I Interferon. Plos One 6, e25858.

35. Ryan, K.A., Bewley, K.R., Fotheringham, S.A., Slack, G.S., Brown, P., Hall, Y., Wand, N.I., Marriott, A.C., Cavell, B.E., Tree, J.A., et al. (2021). Dose-dependent response to infection with SARS-CoV-2 in the ferret model and evidence of protective immunity. Nat Commun 12, 81.

36. Conforti, A., Peruzzi, D., Giannetti, P., Biondo, A., Ciliberto, G., Monica, N.L., and Aurisicchio, L. (2009). A Novel Mouse Model for Evaluation and Prediction of HLA-A2-restricted CEA Cancer Vaccine Responses. J Immunother 32, 744–754.

37. Ferrara, F., and Temperton, N. (2018). Pseudotype Neutralization Assays: From laboratory Bench to Data Analysis. Methods Protoc 1, 8.

38. Giannetti, P., Facciabene, A., Monica, N.L., and Aurisicchio, L. (2006). Individual mouse analysis of the cellular immune response to tumor antigens in peripheral blood by intracellular staining for cytokines. J Immunol Methods 316, 84–96.

39. Bénéchet, A.P., Simone, G.D., Lucia, P.D., Cilenti, F., Barbiera, G., Bert, N.L., Fumagalli, V., Lusito, E., Moalli, F., Bianchessi, V., et al. (2019). Dynamics and genomic landscape of CD8+ T cells undergoing hepatic priming. Nature 574, 200–205.

